# SARS-COV-2 NSP5 Antagonizes MHC II Expresion by Subverting Histone Deacetylase 2

**DOI:** 10.1101/2023.02.10.528032

**Authors:** Nima Taefehshokr, Alex Lac, Angela M Vrieze, Brandon H Dickson, Peter N Guo, Catherine Jung, Eoin N Blythe, Corby Fink, Amena Aktar, Jimmy D Dikeakos, Gregory A Dekaban, Bryan Heit

**Affiliations:** Department of Microbiology and Immunology, and the Western Infection, Immunity and Inflammation Centre, The University of Western Ontario, London, Ontario, Canada, N6A 5C1; Robarts Research Institute, London, Ontario, Canada, N6A 3K7

**Keywords:** SARS-CoV-2, NSP5, Major Histocompatibility Complex II, CIITA, Antigen Presentation, IRF3

## Abstract

SARS-CoV-2 interferes with antigen presentation by downregulating MHC II on antigen presenting cells, but the mechanism mediating this process is unelucidated. Herein, analysis of protein and gene expression in human antigen presenting cells reveals that MHC II is downregulated by the SARS-CoV-2 main protease, NSP5. This suppression of MHC II expression occurs via decreased expression of the MHC II regulatory protein CIITA. This downregulation of CIITA is independent of NSP5’s proteolytic activity, but rather, NSP5 delivers HDAC2 to IRF3 at an IRF binding site within the CIITA promoter. Here, HDAC2 deacetylates and inactivates the CIITA promoter. This loss of CIITA expression prevents further expression of MHC II, with this suppression alleviated by ectopic expression of CIITA or knockdown of HDAC2. These results identify a mechanism by which SARS-CoV-2 limits MHC II expression, thereby delaying or weakening the subsequent adaptive immune response.

**Importance:** SARS-CoV-2 alters the expression of many immunoregulatory proteins to limit and delay the host antiviral response, thereby producing a more severe and longer-lasting infection. Preventing and limiting the activation of helper T cells by reducing MHC II expression on antigen presenting cells is one of these strategies, but while this mechanism was identified early in the pandemic, the mechanism allowing SARS-CoV-2 to limit MHC II expression has remained unclear. Herein, we demonstrate that this occurs via a tripartite interaction between viral NSP5 and host HDAC2 and IRF3, where a complex of NSP5 and HDAC2 is recruited to IRF3 bound to the promoter of CIITA—the master regulator of MHC II expression—with the delivery of HDAC2 then mediating the deacetylation of the CIITA promoter and the suppression of MHC II expression.

## Introduction

First identified in December 2019, severe acute respiratory syndrome coronavirus 2 (SARS-CoV-2) rapidly became a leading global cause of morbidity and mortality, with current vaccination and antiviral strategies greatly reducing mortality. This high virulence is due, in part, to multiple mechanisms enabling SARS-CoV-2 to evade and alter host immune responses, thereby delaying viral clearance and prolonging the infection period [1,2]. Humoral immunity against SARS-CoV-2 wanes quickly, allowing for repeat infections, an issue further compounded by the emergence of new SARS-CoV-2 variants [3–5]. Therefore, understanding the immunoevasion mechanisms of SARS-CoV-2 is important to better understand this disease and to develop better targeted treatments and vaccines.

Although some subsets of professional antigen presenting cells (pAPCs) such as monocyte-derived macrophages and dendritic cells (DCs) do not express ACE2—the canonical receptor for SARS-CoV-2-mediated viral entry— these cells can instead be infected via Fc-receptor dependent phagocytosis of antibody-opsonized virions, and potentially through the efferocytosis of SARS-CoV-2 infected apoptotic cells [6–12]. While SARS-CoV-2 is unable to establish a productive infection in macrophages or DCs, viral early genes are expressed in these cells and drive a multi-pronged immunoevasion response. Firstly, the expression of pro-inflammatory cytokines is induced and contributes to the cytokine storm [12]. This cytokine response is typified by high circulating levels of IL-2, IL-6, IL-7, IL-8, IL-10, IP-10, G-CSF, MCP-1, MIP1-α, and TNF-α [13–15]. While this mechanism drives a potent inflammatory response, the cytokine profile is more typical of bacterial infections, and promotes both NK cell exhaustion and reduced NK cell cytotoxicity, thereby producing a non-productive innate immune response that can exacerbate tissue damage [16,17]. Secondly, SARS-CoV-2 suppresses the antiviral interferon (IFN) pathway, reducing the production of type I and type II IFNs. This suppression is driven by ORF6, which sequesters inactive signal transducer and activator of transcription 1 (STAT1) and STAT2 in the cytosol, thereby preventing their nuclear translocation and blocking the primary signalling pathway that initiates antiviral IFN responses [18]. Moreover, membrane protein and non-structural protein 13 further inhibits IFN-I production by degrading TANK-binding kinase 1 [19,20]. Thirdly, infected macrophages and DCs often die, resulting in long-term depletion of some subsets [21]. Fourthly, SARS-CoV-2 directly suppresses antigen presentation on Major Histocompatibility Complex class I (MHC I) through ORF8-mediated redirection of MHC I trafficking to lysosomes where it is degraded [22], and by ORF6-mediated inactivation of the MHC I transcriptional activator (CITA/NLRC5), thereby limiting the killing of SARS-CoV-2 infected cells by CD8^+^ T cells [23]. Finally, infection of alveolar DCs reduces their ability to migrate to draining lymph nodes and suppresses expression of MHC II and the class II transcriptional activator (CIITA) required for MHC II expression [24]. This suppression of MHC II occurs across a range of pAPCs in SARS-CoV-2 infected patients and in cells infected *in vitro* [25–27]. Moreover, some non-professional antigen presenting cells express MHC II in response to infection – including type II alveolar epithelial cells that are a primary target of SARS-CoV-2 infection in the alveolus [28,29]. However, the cellular mechanism which downregulates MHC II in these cells has remained elusive.

MHC II expression is driven by type II interferon signaling [30]. Activation of the IFN-γ receptor complex leads to the structural rearrangements in the receptor complex and activation of Janus kinase 1 and 2 (JAK1/2) tyrosine kinases and phosphorylation and homodimerization of STAT1, which translocates to the nucleus to induce transcription of IFN-γ-inducible genes [31]. Here, STAT1 activates interferon regulatory factor -1 and -3 (IRF1/3), with STAT1 and IRF1/3 then cooperatively inducing expression of CIITA [32,33]. CIITA, via its intrinsic acetyltransferase activity, can then acetylate histones at the MHC II promoter, which decondenses the chromatin to allow access for transcription factors that regulate MHC II expression [34]. Once the chromatin is opened, CIITA and regulatory factor X (RFX) form an enhanceosome complex on the MHC II promoter which recruits and activates additional transcription factors that induce the transcription of MHC II [35]. While inhibition of STAT1 nuclear import by SARS-CoV-2 ORF6 may account for some inhibition of MHC II expression [18], tissue-resident DCs constitutively express significant amounts of CIITA [36], whereas ORF6 expression is limited until 12-16 hours post-infection [37]; more than sufficient time to induce presentation of SARS-CoV-2 antigens following infection. Moreover, IFN-independent mechanisms can drive MHC II expression in macrophages and DCs [38–40]. Therefore, a more direct form of MHC II suppression is likely invoked by SARS-CoV-2. A critical regulator of MHC II expression is histone deacetylase 2 (HDAC2), which suppresses expression of CIITA and MHC II through deacetylation of histones within their promoters [41]. Gordon *et al*. mapped the SARS-CoV-2 protein interactome and identified non-structural protein 5 (NSP5) as an HDAC2 interactor, and defined a putative NSP5 cleavage site near the nuclear localization signal (NLS) of HDAC2 [42]. NSP5—also known as main protease and 3-chymotrypsin like protease—is translated as part of the polyprotein expressed early after viral entry, and cleaves this polyprotein into 11 individual proteins which form the complex that translates full viral RNA and allows for reproduction of the viral genome and the production of mature virions [43]. Through interactions with HDAC2, NSP5 may mediate the epigenetic reprogramming of infected cells, which in pAPCs may include suppression of MHC II expression. Indeed, epigenetic changes are required for SARS-CoV-2 reproduction [44,45], and similar epigenetic reprogramming is known to suppress MHC II expression in Middle East respiratory syndrome–related coronavirus (MERS-CoV) [46]. Therefore, we tested the hypothesis that SARS-CoV-2 NSP5 inhibits MHC II expression through interactions with HDAC2.

## Results

### SARS-CoV-2 NSP5 Downregulates MHC II in Professional Antigen Presenting Cells

SARS-CoV-2 NSP5 is the main viral protease and plays an essential role in viral infection and pathogenesis [47,48]. As main proteases are required for the processing of coronavirus polyproteins [49], deletion NSP5 or inactivation of NSP5 abrogates productive infection by SARS-CoV-2 [43,49,50] To assess the effects of NSP5 on the MHC II antigen presentation system, primary human monocyte-derived DCs (moDCs) were transduced with either empty or NSP5-expressing lentiviral vectors bearing a zsGreen marker. Flow cytometry was used to quantify total surface expression of total MHC II on transduced (zsGreen^+^) moDCs (**Figure 1A-B, S1A-C**), with NSP5 expression reducing the cell surface expression of MHC II to an extent similar to that observed in SARS-CoV-2 patients (30-50% reduction, **Figure 1C-D**) [25–27]. This downregulation was not a general suppression of the MHC II presentation system, as expression of the co-stimulatory molecule CD86 was not affected by NSP5 expression (**Figure 1E)**. Some pathogens such as Human cytomegalovirus reduce their immunogenicity by diverting intracellular trafficking such that MHC II molecules fail to reach the cell surface [51]. To test this possibility, we transduced J774.2 macrophages with NSP5-expressing or empty vectors, labeled the plasma membrane with wheat-germ agglutinin, followed by labeling for total cellular MHC II. Three-dimensional reconstructions of these cells were used to differentiate between cytosolic/vesicular MHC II and cell-surface MHC II, comparing both non-transduced (zsGreen-negative) to transduced (zsGreen-positive) cells in the same samples (**Figure 1F**). Quantitation of these micrographs revealed no changes in the portion of MHC II localized to the cell surface versus intracellular vacuoles (**Figure 1G**) but did identify the same decrease in MHC II expression that was observed with flow cytometry (**Figure 1D,H**), indicating that NSP5 does not affect the trafficking of MHC II to the cell surface, but rather decreases its overall expression. Moreover, this effect was only observed in NSP5-expressing cells but not in neighbouring non-transduced cells, indicating that this effect is cell-intrinsic and not due to NSP5-induced changes in the expression of cytokines or other secreted factors.

**Figure 1:**
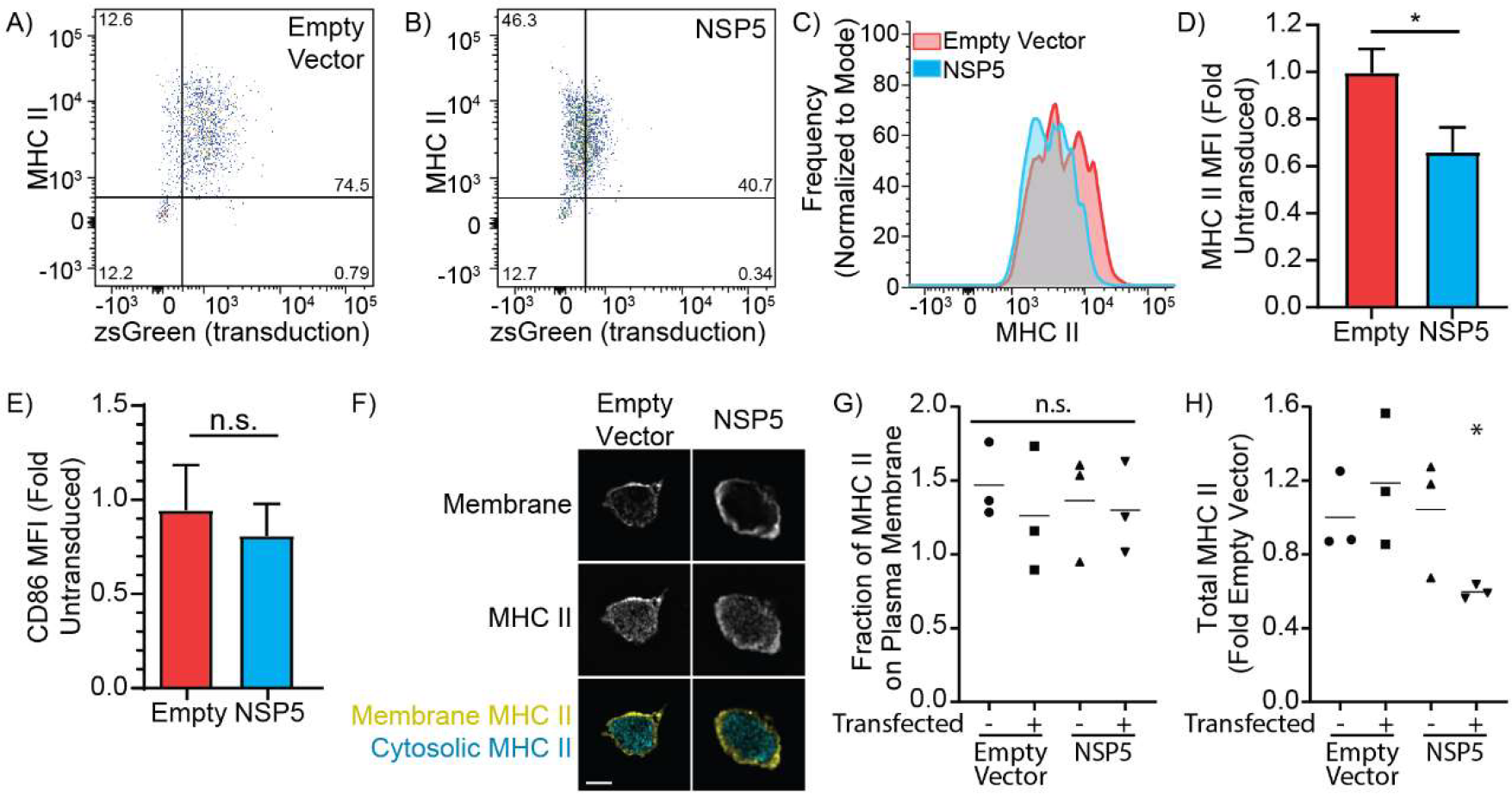
NSP5 Suppresses MHC II in Primary Monocyte-Derived Human Dendritic Cells. moDCs were transduced with lentiviral vectors lacking a transgene (empty vector) or bearing NSP5, with both vectors containing an IRES-zsGreen marker. **A-B)** Representative flow cytometry dot plots showing cell surface MHC II expression and transduction (zsGreen^+^) of moDCs transduced with an empty vector control (A) or NSP5-expressing lentiviral vector (B). **C)** Histogram of cell surface MHC II expression levels on moDCs transduced with an empty vector (red) or NSP5-expressing vector (cyan). **D-E)** Quantification of cell surface MHC II (D) and CD86 (E) in moDCs transduced with either an empty (Empty) or NSP5-expressing (NSP5) lentiviral vector. MFI is normalized to the MFI of the untransduced cells in the Empty-vector condition. **F)** Z-slice through a macrophage stained for MHC II and the plasma membrane, showing the segmentation of vesicular/cytosolic versus surface (membrane) MHC II. Scale bar is 10 μm. **G-H)** Quantification of the fraction of MHC II on the plasma membrane (G) and total cellular MHC II (H) in macrophages that have been transfected either with an empty or NSP5-expressing vector, comparing non-transfected (zsGreen-negative) to transfected (zsGreen-positive) cells in both conditions. Data is representative of, or quantifies a minimum of 3 independent experiments, * = p < 0.05; n.s. = p > 0.05 compared to non-transfected empty vector, Kruskal-Wallace test with Dunn Correction.

### NSP5 Suppresses CIITA and MHC II Transcription

Next, the subcellular localization of NSP5 was determined to identify potential mechanisms accounting for the downregulation of MHC II. Quantitative microscopy of HeLa cells expressing NSP5-FLAG, the ER marker KDEL-GFP, the Golgi marker GalT-mCherry, and with the nuclei stained with Hoechst, determined that approximately half of the cellular NSP5 was localized to the nucleus, with the remainder associated with the ER (**Figure 2A-B**). The nuclear localization of NSP5 was further confirmed by pharmacologically blocking importin-mediated nuclear transport with ivermectin (**Figure 2C-D**). While localization to the ER is consistent with the known role of NSP5 in forming the viral replication complex [52], the role for nuclear NSP5 remains unclear. Unexpectedly, bioinformatic analysis of NSP5 failed to identify either a classical or bipartite nuclear localization signal, or a nuclear export signal. This suggests that NSP5 may be carried into the nucleus through interactions with other cellular proteins, similar to the hepatitis delta antigen [53].

**Figure 2:**
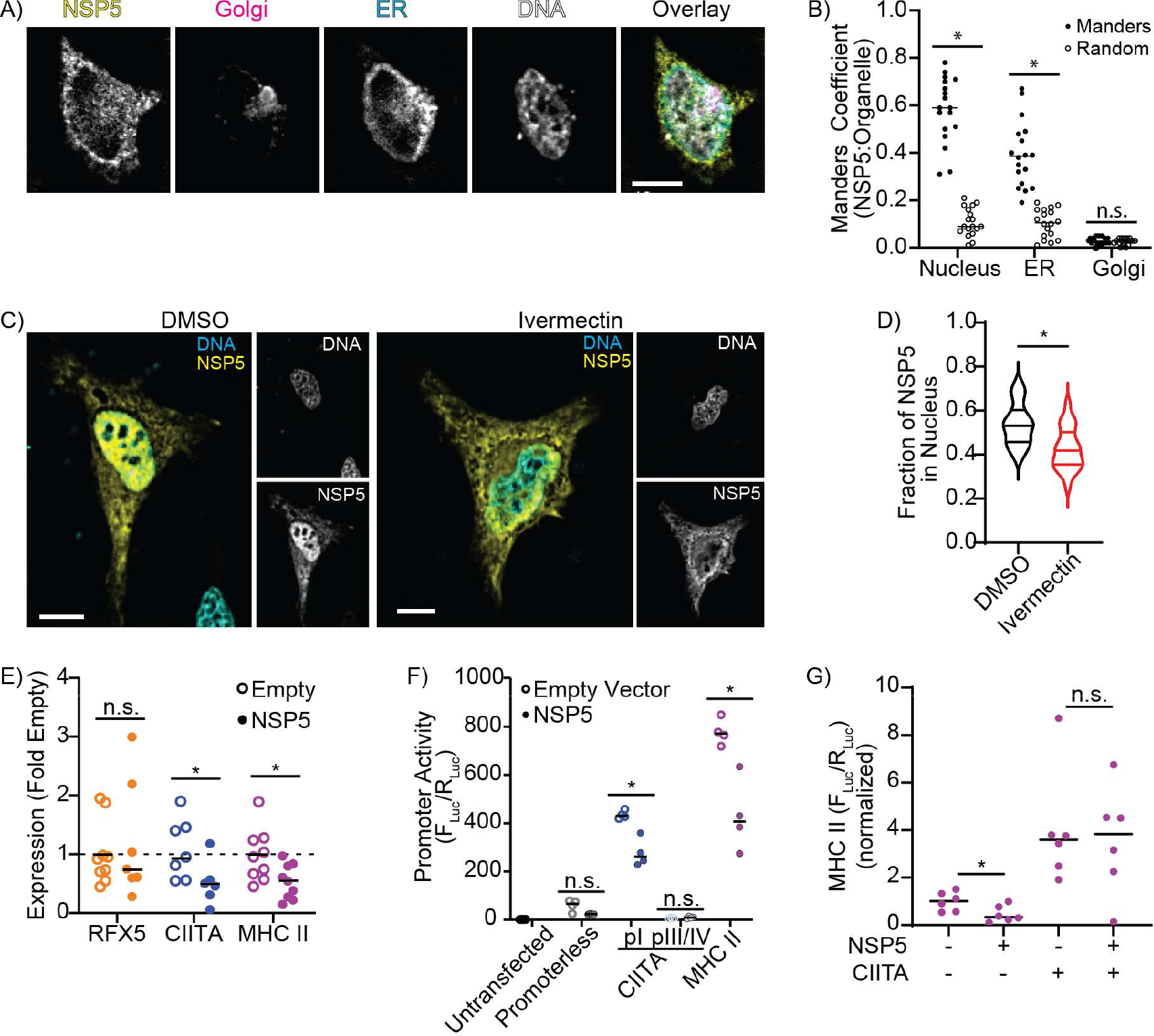
NSP5 Suppresses CIITA and MHC II expression. **A-B)** Fluorescent z-projection (A) and Manders colocalization analysis (B) quantifying the portion of NSP5 colocalized with the nucleus of HeLa cells co-transfected with NSP5-FLAG (yellow), GalT-mCherry (Golgi/magenta), KDEL-eGFP (ER/cyan), and stained with Hoechst (DNA/grey). Manders colocalization analysis compares the fraction of NSP5 colocalized with the nucleus, ER, and Golgi (Manders) to the Manders ratio from the same image when the NSP5 image was randomized (Random). **C-D)** Fluorescent z-projections (C) and quantification (D) of the fraction of NSP5 in the nucleus of vehicle-treated (DMSO) versus ivermectin-treated HeLa cells expressing NSP5-FLAG (yellow) and stained for DNA with Hoechst (cyan). **E)** RT-qPCR quantification of RFX5, CIITA, and MHC II mRNA levels in moDCs transduced with empty or NSP5-expressing lentiviral vectors. **F)** Quantification of the promoter activity of the CIITA pI, CIITA pIII/IV, and MHC II promoters using a dual-luciferase assay in RAW 264.7 macrophages co-transfected with empty or NSP5-expressing lentiviral vectors. **G)** Quantification of MHC II promoter activity in RAW 264.7 macrophages co-transfected with CIITA and NSP5. (-) indicates the sample was transfected with empty vector rather than CIITA or NSP5. F_luc_/R_luc_ was normalized to that of cells transfected with an empty vector. Scale bars are 10 μm. Images are representative of a minimum of 30 cells captured across 3 independent experiments. n = minimum of 3. * = p < 0.05; n.s. = p > 0.05, compared to Random (B), DMSO (D), or Empty Vector (E-F), paired t-test (B) or Mann-Whitney test (D, E, F).

The nuclear localization of NSP5 suggests that downregulation of MHC II occurs via a transcriptional mechanism. To test this hypothesis, we used RT-qPCR to measure the mRNA levels of MHC II, as well as RFX5 and CIITA— two transcription factors which act as master regulators of MHC II expression. Interestingly, while RFX5 expression was unchanged, NSP5 significantly downregulated the expression of CIITA and MHC II relative to the levels normally found on resting moDCs (**Figure 2E**). In humans, CIITA is transcribed from three separate promoters: pI—which drives expression in myeloid cells, pIII—which drives expression in lymphoid cells, and pIV—which drives IFN-γ-induced expression in non-immune cells such as epithelia [54,55]. To assay the activity of the MHC II and CIITA promoters, we constructed dual-luciferase reporters of the MHC II promoter, the CIITA pI promoter, and the CIITA pIII/pIV promoters, followed by quantification of NSP5’s effect on these promoters’ activity in macrophages (**Figure S1D)**. NSP5 expression strongly suppressed both the MHC II promoter and the CIITA pI promoter, whereas the CIITA pIII/IV promoters were minimally active, producing insufficient signal to observe any effect of NSP5 (**Figure 2F**). MHC II transcription is dependent on CIITA, therefore this suppression of MHC II expression may be due to NSP5-dependent repression of CIITA expression, or alternatively, may be a product of NSP5 suppression of both the CIITA and MHC II promoters. To differentiate between these possibilities, we quantified MHC II promoter activity in cells ectopically expressing CIITA and NSP5 (**Figure 2G**). CIITA expression greatly increased MHC II promoter activity, with co-expression of NSP5 having no effect on MHC II promoter activity in the presence of ectopically expressed CIITA. Likewise, we infected A549 cells with SARS-CoV-2—which typically induce MHC II expression in response to infection [56]—but did not observe an upregulation of MHC II or CIITA (**Figure S1E**). These data indicate that NSP5 likely functions by suppressing the transcription of CIITA, with the resulting absence of CIITA then limiting MHC II expression.

### NSP5-Mediated Suppression of MHC II Expression is Dependent on HDAC2

MHC II expression is regulated by CIITA through several mechanisms. Induction of MHC II expression begins with CIITA binding to distal enhancers located several kilobases 5′ to the MHC II promoter [57], followed by acetylation of histones and transcription factors within the core MHC II promoter protein complex by CIITA. This acetylation induces the formation of an enhanceosome complex comprised of CIITA, RFX5, CREB and NF-Y, wherein CIITA activates TAF family transcription factors via its intrinsic protein acetylation and kinase activity, thus initiating MHC II transcription [58,59]. CIITA expression is induced by IFN-γ through the transcription factor IRF1, and by Toll-like receptor (TLR) and IL-1 family cytokines via the transcription factor IRF3 [32,60,61], and is negatively regulated by HDAC2-mediated promoter deacetylation [62]. Critically, SARS-CoV-2 NSP5 has been shown to engage in a non-proteolytic interaction with HDAC2, suggesting that NSP5 utilizes intact HDAC2 to silence the CIITA promoter [42]. Consistent with this model, endogenous HDAC2 co-immunoprecipitated with FLAG-tagged NSP5 in an anti-FLAG immunoprecipitation (**Figure 3A**). As NSP5 is a protease, and proteolytic cleavage of some HDACs are known to promote their activity [63], we immunoblotted for endogenous HDAC2 in cells expressing NSP5, but did not detect the 43 kDa cleavage fragment that would result from NSP5 proteolysis (**Figure 3B**). We confirmed that our ectopically expressed NSP5 was proteolytically active using both a FRET reporter and immunoblotting (**Figure S2**), and using an HDAC2 intramolecular cleavage probe (**Figure S3A-C**), we confirmed that HDAC2 was not cleaved in cells co-expressing NSP5. Knockdown of HDAC2 (**Figure S3D**) had a profound restorative effect on CIITA and MHC II mRNA levels, with HDAC2 knockdown not only reversing – but increasing above baseline – expression of both genes (**Figure 3C**). NSP5 is comprised of a proteolytic domain formed by the interface of the globular A and B domains, which positions two key catalytic residues (H41 and C145) in a binding cleft formed between the two domains. The C-terminal B’ chain folds over this cleft, coordinating both the substrate and three water molecules within the active site [64]. We inactivated the catalytic site by generating NSP5^H41A^ and NSP5^C145S^ point mutants and deleted the proteolytic (NSP5^Δ1-192^) and B’ domains (NSP5^Δ199-306^, **Figures 3D**). The deletion mutants were unstable, with a half-life less than a quarter of that of wild-type or point-mutant NSP5 (**Figure 3E, S3E-F)** and therefore could not be expressed at levels amenable for further experimentation. As expected, both point mutants lost their proteolytic activity (**Figure S2D**,**E**), with immunoprecipitation of the point mutants revealing that both retained some binding capacity for HDAC2 (**Figure 3F**) and retained their ability to suppress CIITA and MHC II promoter activity (**Figure 3G**).

**Figure 3:**
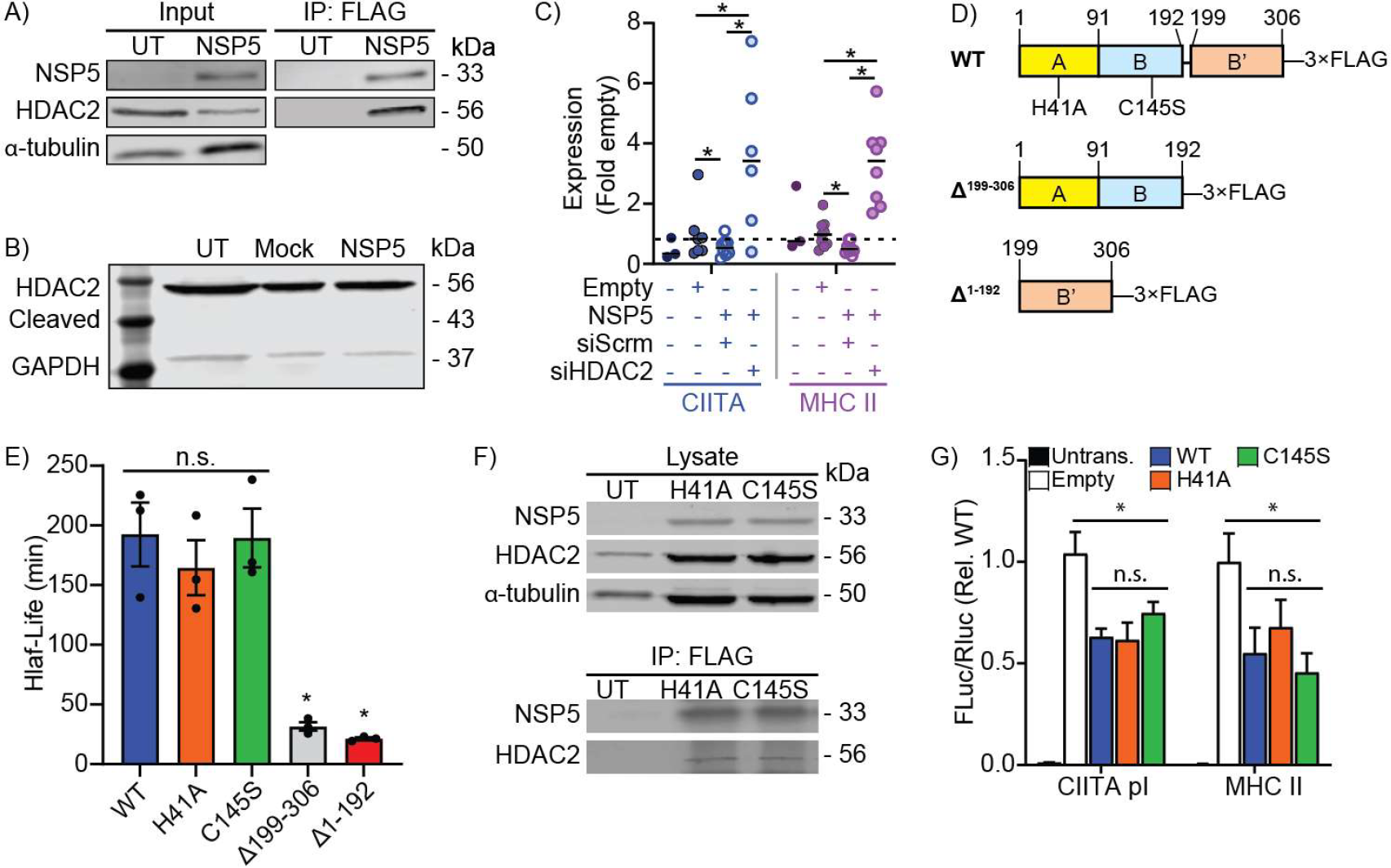
NSP5 Requires HDAC2 to Suppress MHC II Expression. **A)** Co-immunoprecipitation of HDAC2 with NSP5-FLAG from primary human moDCs transduced with empty or NSP5-FLAG-expressing lentiviral vectors. **B)** Absence of detectable cleavage of endogenous HDAC2 in cells ectopically expressing NSP5. Cells are either are untreated (UT), transfected with an empty vector (Mock), or transfected with NSP5, and the blots stained for endogenous HDAC2 and GAPDH. The expected size of NSP5-cleaved HDAC2 is 43.7 kDa. **C)** RT-qPCR quantification of the impact of scrambled (siScrm) or HDAC2-targeting (siHDAC2) siRNA on CIITA and MHC II mRNA levels in primary human moDCs transduced with empty or NSP5-expressing lentiviral vectors. **D)** Protease-inactivating (H41A and C145S) point-mutants, and deletion of the catalytic (A/B, Δ1-192) and ligand-stabilizing (B’, Δ199-306) domains were generated to assay the roles of these sites in NSP5 activity. **E)** Half-life of NSP5 and the H41A, C145S, Δ1-192, and Δ199-306 mutants, as quantified by NSP5 densitometry in cycloheximide-treated cells. **F)** Immunoprecipitation of HDAC2 with NSP5^H41A^ and NSP5^C145S^ mutants. **G)** Dual-luciferase assay quantification of CIITA pI and MHC II promoter activity in RAW264.7 macrophages that were either untransfected (Untrans.), or co-transfected with the CIITA or MHC II luciferase constructs plus either the empty vector (Empty), or with one of wild-type NSP5 (WT), NSP5^H41A^, or NSP5^C145S^ vectors. Data is normalized to Empty. n = 3-5, * = p < 0.05; n.s. = p > 0.05, Kruskal-Wallis test with Dunn correction.

### NSP5 Induces Deacetylation of the CIITA and MHC II Promoters

Given the dependence of NSP5 on HDAC2 for its suppressive effect on MHC II transcription, it is likely that NSP5 is modulating protein acetylation at the CIITA or MHC II promoters. Conventional ChIP was not possible as the large portion of untransduced cells did not allow for accurate quantification of NSP5-driven changes in promoter acetylation. As such, we took advantage of the 30-50% transduction efficiency to compare MHC II and CIITA promoter acetylation between neighbouring transduced versus non-transduced primary human macrophages using FISH-FRET microscopy. This assay measures the transfer of excitation energy from a Cy3-labeled anti-acetyl-lysine antibody to ATTO647-N labeled FISH probes specific to the MHC II promoter plus the 5′ enhancer region, the CIITA-pI promoter plus the 5′ enhancer region, or to the region containing the CIITA pIII and pIV promoters (CIITA pIII/IV, **Figure S1D**), with the zsGreen marker of the lentiviral vector used to differentiate between non-transduced and NSP5-transduced macrophages. In addition to allowing for direct comparisons between NSP5-expressing and non-expressing macrophages sharing an otherwise identical environment, this approach has the additional advantage that both histone acetylation and acetylation of promoter-bound transcription factors are measured, with acetylation of transcription factors such as CIITA known to enhance their activity [65]. Primary human macrophages were transduced with empty or NSP5-expressing vectors and treated with either scrambled or HDAC2-targeting siRNA. Acetylated-lysine staining was concentrated in the nucleus, with weaker staining present in the cytosol, with one or two FISH probes in-focus within each nucleus (**Figure 4A**). Neither NSP5 expression, nor HDAC2 depletion, altered the quantity or distribution of cellular acetyl-lysine staining (**Figure 4B-C**), indicating that neither NSP5 expression nor HDAC2 knockdown globally affected lysine acetylation. In untransduced cells, and in cells transduced with an empty vector, a significant FRET signal could be observed at the CIITA pI and MHC II promoters, while a much weaker FRET signal was detected at the CIITA pIII/IV promoter (**Figure 4D-F**), consistent with the pI promoter driving CIITA expression in myeloid cells. Critically, ectopic expression of NSP5 significantly reduced acetylation at all three promoters, with HDAC2 knockdown restoring promoter acetylation in NSP5-expressing cells (**Figure 4D-F**).

**Figure 4:**
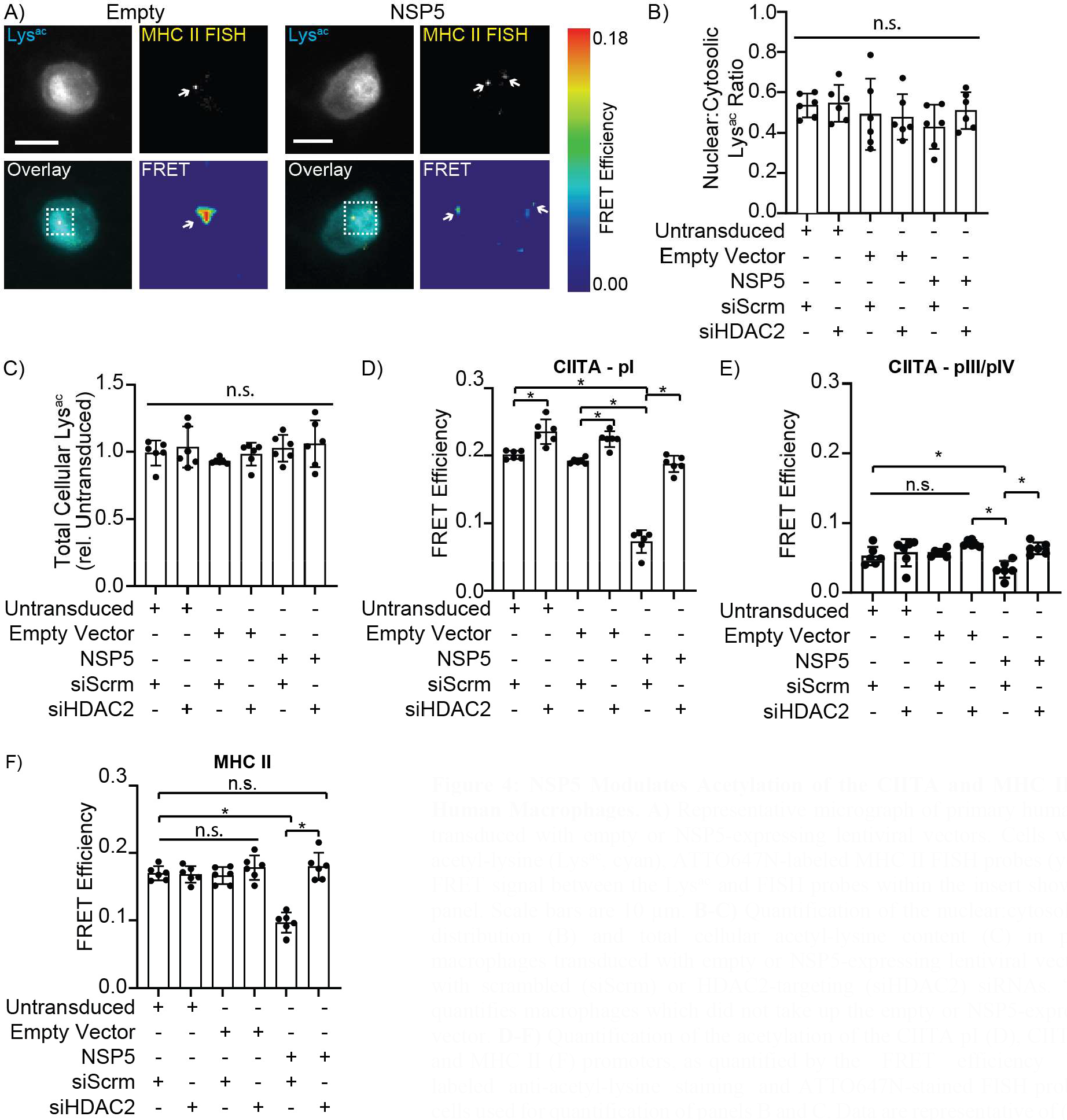
NSP5 Modulates Acetylation of the CIITA and MHC II promoters in Human Macrophages. **A)** Representative micrograph of primary human macrophages transduced with empty or NSP5-expressing lentiviral vectors. Cells were stained for acetyl-lysine (Lys^ac^, cyan), ATTO647N-labeled MHC II FISH probes (yellow), with the FRET signal between the Lys^ac^ and FISH probes within the insert shown in the FRET panel. Scale bars are 10 μm. **B-C)** Quantification of the nuclear:cytosolic acetyl-lysine distribution (B) and total cellular acetyl-lysine content (C) in primary human macrophages transduced with empty or NSP5-expressing lentiviral vectors and treated with scrambled (siScrm) or HDAC2-targeting (siHDAC2) siRNAs. “Untransduced” quantifies macrophages which did not take up the empty or NSP5-expressing lentiviral vector. **D-F)** Quantification of the acetylation of the CIITA pI (D), CIITA pIII/pIV (E), and MHC II (F) promoters, as quantified by the FRET efficiency between Cy3-labeled anti-acetyl-lysine staining and ATTO647N-stained FISH probes in the same cells used for quantification of panels B and C. Data are representative of (A) or quantifies (B-F) three independent experiments. * = p < 0.05; n.s. = p > 0.05, Kruskal-Wallis test with Dunn correction. Flat bars indicate the statistical significance for all groups beneath the bar; legged bars indicate the statistical significance between the groups below the legs.

### NSP5 Targets the CIITA Promoter via Interactions with IRF3

We next quantified the subcellular distribution of HDAC2 and the NSP5 mutants, and found that inactivation of the NSP5 catalytic site had no impact on the distribution of either protein (**Figure 5A-B**). Although poorly expressed, the rare cells expressing NSP5 deletion mutants lacking the proteolytic (NSP5^Δ1-192^) or B′ (NSP5^Δ199-306^) domains displayed a hyper-nuclear localization compared to wild-type, but also had no impact on the distribution of HDAC2, indicating that HDAC2 is unlikely to be the protein used by NSP5 to gain access to the nucleus (**Figure 5A-B**). Naik *et al*. demonstrated that NSP5 can interact with IRF3, and IRF3 is known to bind to the CIITA PI promoter, suggesting that NSP5 may deliver HDAC2 to the CIITA promoter via interactions with IRF3 [66–68]. Consistent with this study, we found that IRF3 co-precipitated with NSP5, and that this interaction was maintained following inactivation of the NSP5 catalytic site (**Figure 5C**). Then, using FISH-FRET analysis in A549 cells, we determined that IFN-γ stimulation increased CIITA PI, PIII/IV and MHC II promoter acetylation, and as expected, NSP5 expression inhibited acetylation of all three promoters (**Figures 5D-F, S4)**. Critically, this suppression of promoter acetylation by NSP5 was largely reversed by an IRF3-targeting siRNA, consistent with a model wherein NSP5 delivers HDAC2 to the CIITA promoter via interactions with IRF3 (**Figures 5D-F, S4**). To confirm the specificity of this interaction we generated a PIII/PIV promoter luciferase construct lacking the putative IRF binding site (**Figure S1D**). Deletion of this site reduced the activity of the PIII/PIV promoter by ∼30%, but critically, this deletion protected the promoter from further downregulation by NSP5 (**Figure 5G**). These data demonstrate that an interaction between IRF3 and NSP5 is required to deliver NSP5 to the CIITA promoter, where it then mediates the HDAC2-dependent deacetylation and inactivation of CIITA expression.

**Figure 5.**
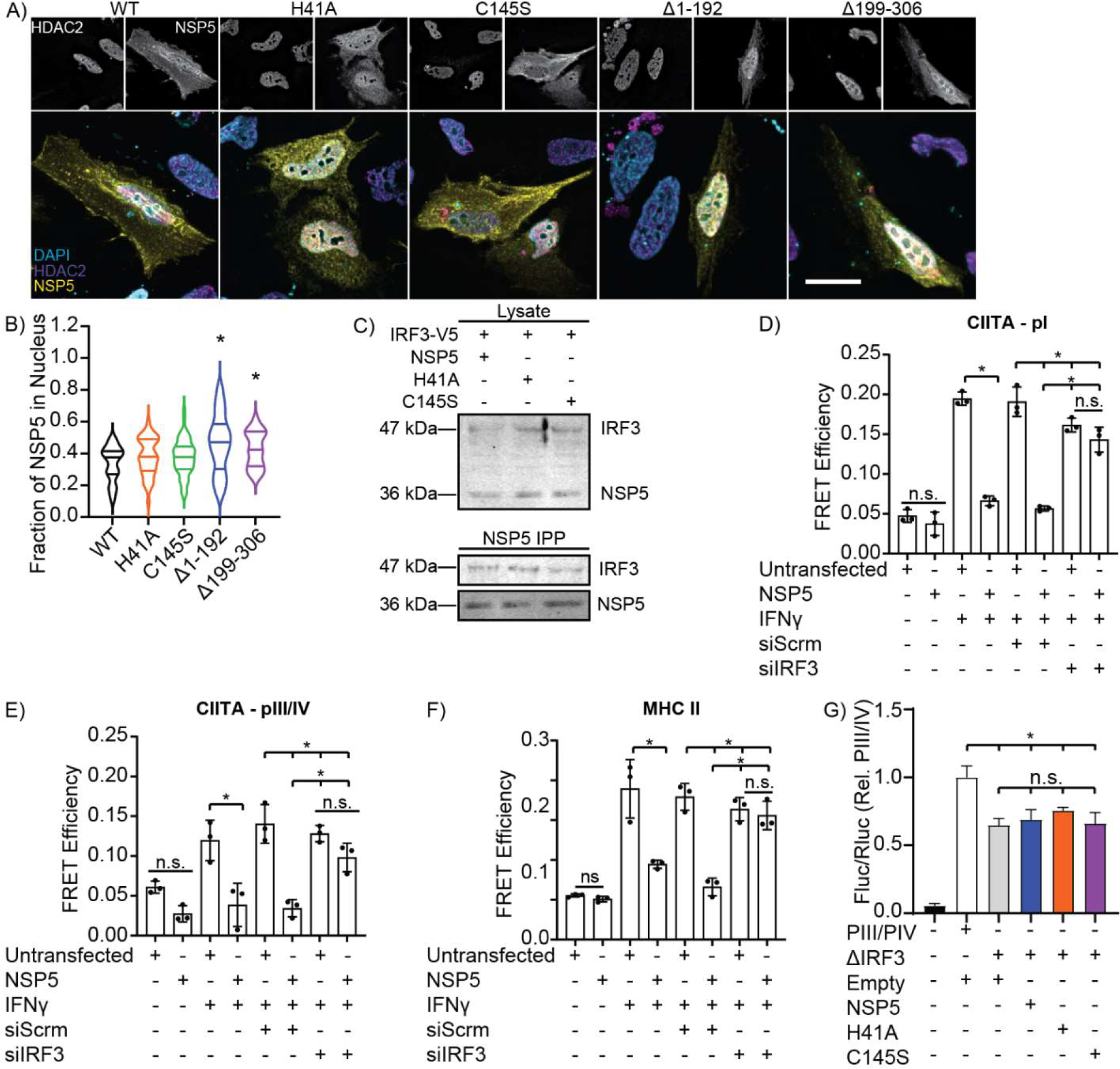
NSP5 Suppresses CIITA Promoter Activity via IRF3. **A-B)** Fluorescent micrographs (A) and quantification (B) of NSP5 (Yellow) and HDAC2 (Magenta) nuclear localization in HeLa cells transfected with wild-type (WT) or mutant NSP5. The nucleus has been stained with DAPI (Cyan). Scale bar is 10 μm. **C)** IRF3 co-immunoprecipitation with wild-type NSP5 and with the H41A and C145S NSP5 catalytic mutants. **D-F)** Quantification of FISH-FRET at the CIITA pI (D), CIITA pII/IV (E), and MHC II (F) promoters in NSP5-transfected A549 cells treated with IFNγ and a non-targeting (Scrm) or IRF3-depleating siRNA. **G)** Impact of wild-type (NSP5) versus protease-inactivated H41A and C145S NSP5 mutants on the activity of the wild-type (PIII/PIV) or IRF3-binding site deleted (ΔIRF3) CIITA PIII/PIV promoter in A549 cells, as quantified by a dual-luciferase assay. Empty = cells are transfected with the empty vector used to express NSP5. Images are representative of a minimum of 30 cells imaged over 3 independent experiments. Data is presented as mean ± SEM (B,H) or quartiles, n = 3, * = p < 0.05; n.s. = p > 0.05 compared to Empty (B), WT (D), or the indicated groups (E-G), Kruskal-Wallis test with Dunn correction.

### IRF3 Targeting is Unique to SARS-CoV-2

It seemed unusual that a virus whose closest known wild ancestor is found in bats (BANAL-20-236) would have activity against human MHC II and CIITA promoters [69,70], however, this may represent a potent immunoevasion activity that could be conserved across *coronaviridae*. Alternatively, NSP5 may have undergone significant evolution following the zoonotic transmission of SARS-CoV-2 to humans, or this activity may be restricted to SARS-CoV-2 and its closely related betacoronaviruses. Phylogenetic and amino acid conservation analysis revealed that HDAC2 is strikingly conserved across the vertebrate clade, with most residues completely conserved between humans and a range of vertebrates known to be infected by SARS-CoV-2 or to frequently contact humans (**Figure 6A**). In marked contrast, IRF3 was poorly conserved across the same vertebrates (**Figure 6B**), and NSP5 was only poorly conserved across the major coronavirus clades (**Figure 6C**). While NSP5 is highly divergent across *coronaviridae*, it is completely conserved within the sarbecovirus sub-clade of the betacoronaviruses, which includes SARS-CoV, SARS-CoV-2, and bat coronavirus BANAL-20-236 (**Figure 6C**). While the highly conserved nature of HDAC2 would make it a good candidate as a pan-species target of NSP5, the higher diversity of IRF3 may limit the extent to which coronaviruses can utilize IRF3 to target NSP5/HDAC2 complexes to specific promoters. To assess these possibilities, we transfected A549 cells with a V5-tagged IRF3 and with a FLAG-tagged NSP5 from SARS-CoV-2, the alphacoronavirus 229E, or HKU1 – a member of the embecovirus sub-clade of the betacoronaviruses. Interestingly, HDAC2 co-immunoprecipitated with NSP5 from all three coronaviruses, whereas IRF3 only co-immunoprecipitated with NSP5 from SARS-CoV-2 (**Figure 6D**). Consistent with this observation, NSP5 from SARS-CoV-2 suppressed promoter activity from the PIII/PIV CIITA promoter, but a similar suppression of CIITA was not observed with NSP5 from either 229E or HKU1 (**Figure 6E**). Combined, these data indicate that NSP5/HDAC2-mediated epigenetic re-programming is potentially conserved across many coronaviruses, while the IRF3-dependent “bridging” of this NSP5-HDAC2 complex to the CIITA promoter is likely limited to the sarbecoviruses.

**Figure 6:**
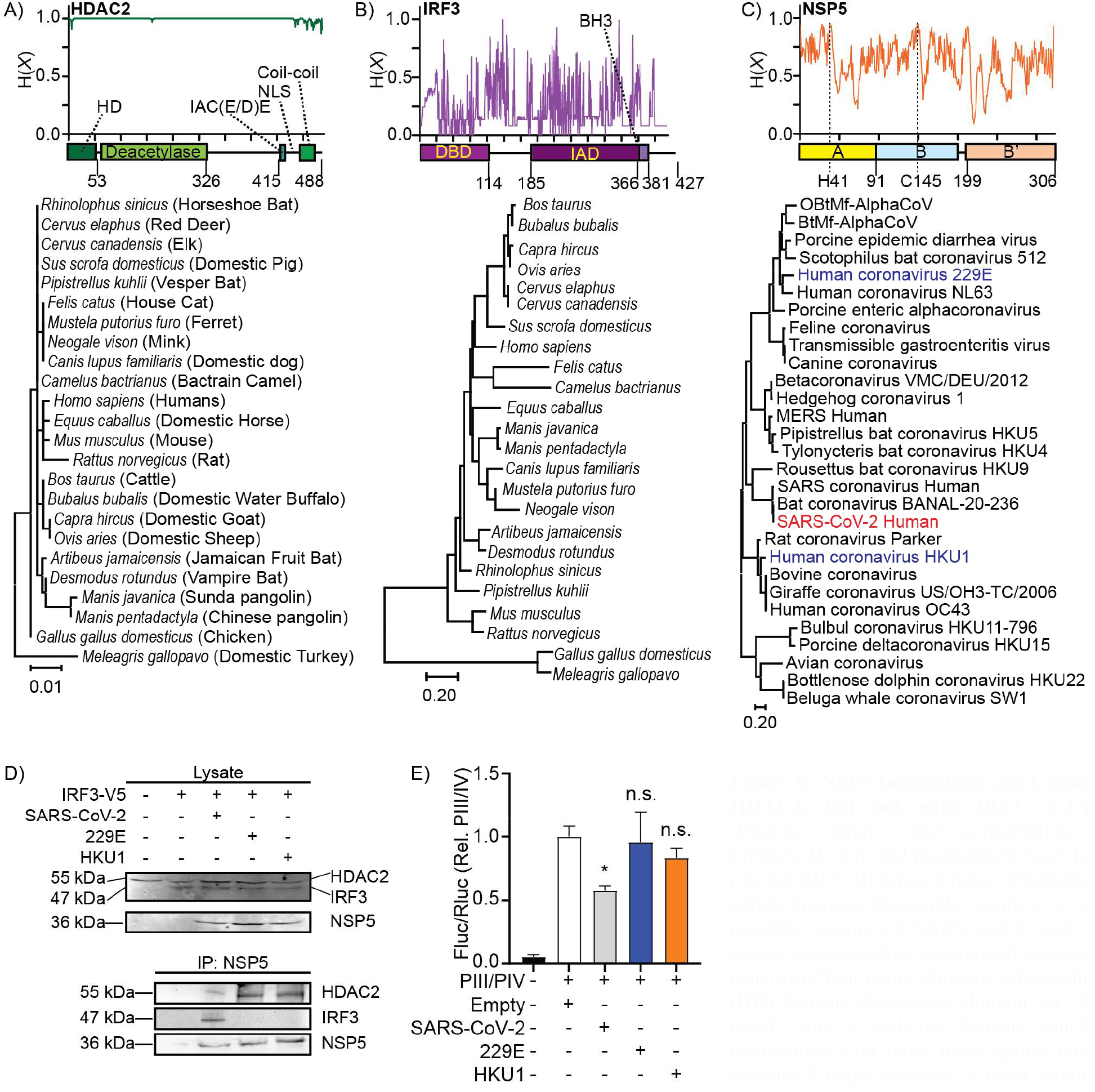
NSP5 Interactions are Conserved with HDAC2, but not with IRF3. **A-C)** Domain structure, amino acid conservation (Shannon Entropy, H (X)), and phylogenetic trees for HDAC2 (A) and IRF3 (B) across a range of vertebrate species which humans frequently contact or which are possible vectors of SARS-CoV2, and NSP5 (C) across representative coronaviral species. HDAC2 consists of four major domains, a homodimerization (HD) domain, deacetylase domain, an IAC (E/D)E motif, and a coil-coil domain which mediate interactions with other transcription factors. IRF3 contains 3 major domains: a DNA binding domain (DBD), an IRF-Association Domain (IAD), and a BH3 domain. NSP5 contains a bipartite protease domain (A/B) with two critical catalytic residues (H41 and C145), and a B’ C-terminal domain which stabilizes ligands in the protease domain. **D)** Co-immunoprecipitation of endogenous HDAC2 from cells co-transfected with IRF3-V5 and with a FLAG-tagged NSP5 from SARS-CoV-2, from Human coronavirus 229E, or from Human coronavirus HKU1. **E)** Dual-luciferase quantification of CIITA promoter PIII/PIV in INF-γ stimulated A549 cells co-transfected NSP5 from either SARS-CoV-2, Human coronavirus 229E, or Human coronavirus HKU1. “Empty” indicates cells co-transfected with the dual-luciferase vectors and the empty version of the vector used to express NSP5. n = 3, * = p < 0.05, n.s. = p > 0.05, compared to empty-vector transduced cells, Kruskal-Wallis test with Dunn correction.

## Discussion

In this study we demonstrate that SARS-CoV-2 NSP5 suppresses expression of CIITA across a range of professional and non-professional antigen presenting cells, thereby suppressing MHC II expression. This suppression of CIITA expression occurs via a pathway in which NSP5 delivers HDAC2 to the CIITA promoter via interactions between NSP5 and promoter-bound IRF3. HDAC2 then deacetylates histones at the CIITA promoter, decreasing CIITA expression, thereby blocking MHC II expression (**Figure S5**). These findings explain the decreased MHC II expression observed in COVID-19 patients and in *in vitro* infection models [25–27]. This suppressive mechanism occurred across multiple types of professional APCs (human moDCs, human macrophages, and a mouse macrophage cell line) and in human non-professional APCs (type II alveolar epithelial cells). As SARS-CoV-2 infects pAPCs including DCs, macrophages, and B cells—as well as non-professional APCs in the lung—this suppression of MHC II has the potential to dramatically decrease the activation of CD4^+^ T cells in COVID-19 patients [6–8]. Such a loss of CD4^+^ T cell activity likely contributes to the shorter-lived humoral immunity and propensity for reinfection of COVID-19, and may contribute to the aberrant formation of memory and effector CD4^+^ T cell subsets in individuals with severe disease [24,71–74].

The suppression of MHC I and MHC II expression, or removal of these proteins from the cell surface, is a commonly employed viral immunoevasion strategy. For MHC I, mis-directing or re-internalizing MHC I after its translation are the predominant mechanisms used to limit immunogenicity of infected cells [75]. For example, HIV-1 Nef mediates the AP-1-dependent endocytosis of MHC I and its sequestration in a Golgi-proximal compartment, thus limiting cytotoxic T cell activity against infected cells [76,77]. Similarly, SARS-CoV-2 utilizes ORF8 to direct MHC I into lysosomes for degradation, and uses ORF6 to suppress MHC I expression by downregulating the transcription factor CITA/NLRC5 [18,22]. Influenza A and B also downregulate surface MHC I via proteasomal degradation or via endocytosis and sequestration in cytosolic vacuoles, respectively [78]. Downregulation of MHC II is rarer, as only viruses that infect APCs have the potential to exhibit this activity. HSV-1 reduces cell surface levels of MHC II by directing MHC II into multivesicular bodies [79,80]. HIV-1 Nef, through accumulation of αβIi complexes in intracellular vesicles, suppresses trafficking of peptide-loaded MHC II to the cell surface [51]. Human cytomegalovirus protein US2 degrades HLA-DR-α and DM-α chains, while the US3 protein competes with the invariant chain for binding to MHC II α/β complexes, thereby restricting MHC II intracellular trafficking [81,82]. Transcriptional downregulation of MHC II is also observed, often through targeting of CIITA. Similar to what we have reported in this study, Kaposi sarcoma-associated herpesvirus targets the IRF3 binding site in the CIITA promoter—but unlike SARS-CoV-2—this virus does so through expressing a viral IRF3 that inhibits CIITA expression [67]. Epstein-Barr virus impairs CIITA expression in B cells by suppressing the activity of E47 and PU.1, thereby preventing these transcription factors from binding to the pIII promoter [83,84]. The extent to which coronaviridae can suppress MHC II is unclear. We found that the NSP5/HDAC2/IRF3-dependent suppression of MHC II was restricted to the betacoronavirus SARS-CoV-2—but given that NSP5 from SARS-CoV and BANAL-20-236 share 100% homology with SARS-CoV-2 NSP5, this suppression of MHC II expression is expected to be conserved across the sarbecoviruses. This activity was not found in the more distally related betacoronavirus HKU1, nor was it found in the alphacoronavirus 229E. Other members of the coronaviridae appear to have independently evolved mechanisms to suppress MHC II expression. The betacoronavirus MERS-CoV also transcriptionally downregulates MHC I and MHC II, but unlike SARS-CoV-2, does so in the presence of elevated CIITA expression [46]. The alphacoronavirus 229E does not suppress antigen presentation, and instead kills infected dendritic cells before they can complete maturation and begin presenting antigens [85]. As most tested coronaviruses fail to infect pAPCs [86–88], it may be that suppression of MHC II arises independently in those coronaviruses that evolved to infect pAPCs, which would account for the lack of conservation of MHC II suppression mechanisms across the coronaviridae.

The proteolytic activity of coronaviral NSP5 is known to be important for its immunosuppressive activity. Porcine deltacoronavirus NSP5 suppresses host antiviral IFN signaling by proteolysis of the host antiviral proteins NF-kappa-B essential modulator [89], STAT2 [90], and mRNA decapping protein 1a [91]. Previous computational studies also predicted a SARS-CoV-2 NSP5 cleavage site (^380^VQMQ|AIPE^387^) in HDAC2, with cleavage removing an 11.6 kDa C-terminal fragment containing the HDAC2 NLS [42]. This cleavage was proposed to reduce HDAC2 localization to the nucleus and thus limit its ability to attenuate inflammatory responses. Surprisingly, we did not observe evidence that NSP5 cleaves HDAC2, with no cleaved HDAC2 (∼44 kDa) appearing in our immunoblotting experiments, nor was cleavage detected in our intramolecular FRET assays. This is consistent with the work of Naik *et al*. who observed a similar non-proteolytic interaction of NSP5 with HDAC2 and IRF3, although in contrast, herein we observed a suppressive effect of this interaction on MHC II expression while Naik *et al*. found that this interaction was dispensable for the suppression of IL-6, IL-1β, and IFN-β [66]. Further demonstrating that cleavage is not required for this interaction, the catalytically inactive NSP5^H41A^ and NSP5^C145S^ point mutants maintained their interaction with HDAC2 and IRF3, and had the same suppressive effect on CIITA and MHC II promoter activity as wild-type NSP5. Why HDAC2 is not cleaved by NSP5, despite containing an accessible SARS-CoV-2 NSP5 consensus sequence, is unclear. However, non-proteolytic functions of SARS-CoV-2 NSP5 have been reported, including inducing SUMOylation of MAVS to promote inflammation [92] and suppressing activation of RIG-I by preventing formation of antiviral stress granules via interactions with G3BP1 [93]. Additionally, coronavirus-mediated epigenetic reprogramming has been reported previously, with both SARS-CoV and MERS-CoV engaging in targeted alterations of inhibitory and activating histone modifications across a range of genes [94]. Consistent with this observation, we found that SARS-CoV-2 NSP5 selectively inhibited expression of MHC II and CIITA without affecting expression of CD86 and RFX5, or globally suppressing acetylation, with this targeting driven by the selective delivery of NSP5 to the CIITA promoter via interactions with promoter-binding IRF3.

HDAC2 is a promising target for treatment of SARS-CoV-2 infection. Indeed, epigenetic changes consistent with HDAC2 activity drives the cytokine storm that is responsible for much of the pathology seen in SARS-CoV-2 patients, with the degree of epigenetic change correlated to disease severity [44,95–97]. NSP5 is thought to drive much of the cytokine response in the infected lung via HDAC2, and indeed, HDAC2 is required for the upregulation of many pro-inflammatory molecules in endothelial and myeloid cells undergoing SARS-CoV-2 infection [98–100]. HDAC inhibitors exhibit potent anti-inflammatory effects and therefore may antagonize many of the inflammatory pathways activated by SARS-CoV-2 [101,102]. Outside of inflammation, HDACs positively regulate ACE2 expression, and consequentially, HDAC2 inhibitors decrease SARS-CoV-2 entry and replication through reducing viral entry [101,103,104]. Moreover, HDAC inhibitors downregulate pro-inflammatory cytokines, reduce lung fibrosis, prevent viral entry into the central nervous system, and decrease neurological damage [102,105,106]. Thus, HDAC2 inhibition may improve patient outcomes through multiple mechanisms in addition to restoration of MHC II expression. Critically, targeting a host protein would reduce the likelihood of SARS-CoV-2 evolving resistance to this treatment.

Understanding how SARS-CoV-2 modulates the MHC II antigen presentation pathway provides an important insight into the immunoevasion tactics used by this virus and may help to provide directions for the design of future COVID-19 vaccines or therapeutics. This study identifies one mechanism through which SARS-CoV-2 suppresses MHC II expression. Furthermore, our data indicate that SARS-CoV-2 may utilize NSP5 to modulate a broad array of immune responses via targeting HDAC2, which in addition to its effects on MHC II expression identified herein, has also been identified as a positive regulator of the cytokine storm that underlies much of the pathology of COVID-19. Indeed, HDAC2 inhibition has been proposed as a therapeutic approach for COVID-19, with the findings from this study further validating HDAC2 inhibitors as potentially valuable treatments for this disease [105].

## Materials & Methods

### Materials

The TGN46-GFP and KDEL-mRFP plasmids were gifts from Dr. Sergio Grinstein (Hospital for Sick Children, Toronto, Canada). A549 ACE2 TMPRSS2 cells were a gift from Matthew Miller (McMaster University, Hamilton, Canada). The pMD2.G (plasmid 12259) and pDR8.2 (plasmid 12263) packaging vectors, pcDNA3-myc-CIITA (plasmid 14650), and human IRF3-V5 (plasmid 32713) were purchased from AddGene. All DNA primers and synthesized genes were from IDT (Coralville, Iowa), and the sequences for all primers used in this study can be found in Supplemental Table S1. Tissue culture medium, fetal bovine serum (FBS), and trypsin were from Wisent (St. Bruno, Canada). Recombinant cytokines were from Peprotech (Cranbury, NJ). All cell lines and Lympholyte-poly were from Cedarlane labs (Burlington, Canada). Polybrene, ivermectin, and 100K Amicon centrifugal filters were purchased from EMD Millipore Corp (USA). CD14 Positive Cell Selection Kit, FcBlock, and anti-DYKDDDDK Tag (L5) were from BioLegend (San Diego, California). The #1.5 thickness coverslips and 16% paraformaldehyde (PFA) were from Electron Microscopy Sciences (Hatfield, PA). FDA-traceable PLA filament was purchased from Filaments.ca (Mississauga, Canada). RNeasy Mini Kit was from Qiagen (Germantown, MD). Permafluor, versene, WGA-Alexa Fluor 647, DAPI, Hoescht, HALT protease/phosphatase inhibitors, Dithiobis[succinimidyl propionate], hygromycin, and dithio-bismaleimidoethane, were purchased from ThermoFisher Canada (Mississauga, Canada). The suppliers and all antibodies and labeling reagents used in this study can be found in Supplemental Table S2. Atto647N NT Labeling Kit was from Jena Biotech (Jena, Germany). Accell cell-penetrating SMARTpool scrambled and HDAC2-targeting siRNAs were from Horizon Discovery (Cambridge, UK). Instagene, iScript Select cDNA Synthesis Kit, 4%-20% SDS-PAGE gels, SsoFast EvaGreen Supermix, and all protein blotting reagents/gels were from BioRad Canada (Mississauga, Canada). Renilla luciferase internal control vector pRL-TK and the dual luciferase reporter assay kit were from Promega (Madison, WI). Phusion PCR enzyme, all restriction enzymes, HiFi Gibson Assembly Kit, and T4 DNA ligase were from NEB Canada (Whitby, Canada). The pLVX-zsGreen lentiviral vector and Retro-X Universal Packaging System were purchased from Takara Bio (San Jose, California). All lab plasticware, PolyJet and GenJet transfection reagent, and DNA isolation kits were from FroggaBio (Concord, Canada), and all laboratory chemicals were from Bioshop Canada (Burlington, Canada).

### Cloning and Retroviral Packaging

The NSP5 RNA sequence from the Wuhan SARS-CoV-2 strain [107], Human Coronavirus 229E, and Human Coronavirus HKU1 were synthesized such that a start and stop codon were added to the 5’ and 3’ end of the NSP5 sequence, along with 20 bp of homology to the pLVX-zsGreen vector at the EcoRI restriction site. The resulting NSP5 sequences were cloned into EcoRI-digested pLVX-zsGreen by Gibson assembly. Point mutants were generated by amplifying the entirety of this original vector with phosphorylated primers that incorporate the point mutation in the first base pair of the forward primer, while deletion mutants were generated by amplifying the vector from either side of the desired deletion with phosphorylated primers. After amplification, the parental plasmid was removed by DpnI digestion and the amplicons circularized with T4 DNA ligase. All primers used for RT-qPCR can be found in Supplemental Table S1. To produce pseudotyped lentivirus containing empty vector or NSP5, 3×10^6^ HEK293T cells were grown in 75 cm^2^ tissue culture flasks, then transfected with PolyJet transfection reagent (500 μL complex containing 4 μg pMD2.G and 10 μg pDR8.2 packaging vectors, and 10 μg of pLVX expression vector). Following transfection, cells were incubated at 37°C/5% CO_2_ for 18 hr at which point the media was exchanged for 8 mL of DMEM supplemented with 10% FBS and returned to the incubator for 48 hr. The media was transferred to a sterile 50 mL conical centrifuge tube and topped up to 20% FBS, then centrifuged at 4000×g for 5 minutes, and the supernatant filtered with a 0.2 μm syringe filter into a new 50 mL conical tube. The pseudotyped lentivirus was then concentrated using a 100 kDa centrifugal filter unit at 4000×g at 4°C for 45 min per 15 mL of filtrate. Concentrated pseudotyped lentivirus was aliquoted and stored at -80°C and thawed at room temperature prior to use. Dual-luciferase and FISH-FRET

### Human Macrophage and Dendritic Cell Culture, Transduction, and siRNA Treatment

The collection of blood and cells from healthy donors was approved by the Health Science Research Ethics Board of the University of Western Ontario and was performed in accordance with the guidelines of the Tri-Council policy statement on human research. Blood was drawn into heparinized vacuum collection tubes, layered on an equal volume of Lympholyte-poly and centrifuged at 300×g for 35 min at 20 °C. The top band of peripheral blood mononuclear cells was collected and washed once (300×g, 6 min, 20 °C) with phosphate-buffered saline. For dendritic cell differentiation, a CD14 selection kit was used to isolate monocytes according to manufacturer’s instruction. The selected CD14^+^ cells were cultured in RPMI-1640 + 10% FBS and 1% antibiotic–antimycotic solution with GM-CSF (100 ng/ml) and IL-4 (100 ng/ml) for 4 days to yield immature monocyte-derived DCs (moDCs). To produce macrophages, selected CD14^+^ cells were cultured in the presence of M-CSF (10 ng/ml) for 6 days. ∼1 × 10^6^ moDCs or macrophages were centrifuged with 20 transducing units of lentiviral vectors per cell at 800×g at 32°C for 90 min with Polybrene (10 μg/ml). After centrifugation, the cells were incubated at 37°C with 5% CO_2_, and 8 hr later fresh media plus cytokines were added and the cells incubated for 72 hr. For siRNA knockdown, 1 μM of Accell cell-permeant siRNA was added to the cells and incubated at 37°C with 5% CO_2_ for 72-96 hr.

### Cell Line Culture and Transfection

HeLa, RAW264.7, and J774.2 cells were cultured in DMEM supplemented with 10% FBS, while A549 cells were cultured in Ham’s F12 supplemented with 10% FBS, and were grown at 37°C/5% CO_2_ incubator. Cells were split 1:10 upon reaching >80% confluency by either scraping cells into suspension (RAW and J774) or by trypsinization, diluting in fresh medium, and replating in a new tissue culture flask. GenJet DNA transfection reagent was used to transfect plasmids into HeLa cells, as per the manufacturer’s instructions. Briefly, for each well in a 6-well plate, 1 μg of DNA was diluted into 50 μL of serum-free DMEM, followed by 3 μL of Genjet reagent. The resulting mixture was incubated for 10 min at room temperature, and then added dropwise to the HeLa cells. Cells were incubated for at least 18 hr at 37 °C in a 5% CO_2_ before collection. A549 cells were transfected using Lipofectamine 3000, transfecting 1 μg of DNA per well of a 12-well plate, using 2 μL of P2000 and 3 μL Lipofectamine per transfection. A Neon transfection system (Thermo Fisher Scientific AG) was used to transfect plasmids into RAW264.7 cells according to the manufacturer’s instructions. Briefly, 1×10^6^ cells were resuspended in 10 μL of buffer R containing 5 μg of plasmid DNA and electroporated using a single 20 ms pulse at 1680V. Lentiviral transductions were used for transducing plasmids into J774.2 cells as described above.

### A549 ACE2 TMPRSS2 Cell SARS-CoV-2 Infection

A549 ACE2 TMPRSS2 cells were cultured in Dulbecco’s Modified Essential Media (DMEM) supplemented with penicillin (100 U/mL), streptomycin (100 mg/mL), HEPES, L-Glutamine (0.3 mg/mL), 10% FBS. ACE2 and TMPRSS2 expression was maintained through 700 μg/mL G418 and 800 μg/mL hygromycin supplementation, and cells were cultured at 37°C, 5% CO_2_ and 100% relative humidity. One day prior to infection, 2×10^4^ A549 ACE2 TMPRSS2 cells were seeded per well of 96 well plate and cultured overnight for cell monolayer to adhere in G418- and hygromycin-deficient DMEM (37°C, 5% CO_2_). On the day of infection, 1×10^4^ TCID50/mL SARS-CoV-2 USA-WA1/2020 virus strain was prepared in MEM supplemented with penicillin (100 U/mL), streptomycin (100 mg/mL), HEPES, L-Glutamine (0.3 mg/mL), 0.12% sodium bicarbonate, 2% FBS and 0.24% BSA in a Biosafety Level 3 laboratory (ImPaKT Facility, Western University). Media was aspirated from 96 well plates and replaced with a volume corresponding to 500 TCID50 virus per well. Uninfected wells received an equivalent volume of MEM + 2% FBS lacking virus. All wells were then incubated for one hour (37°C, 5% CO_2_), at which point virus inoculum or media was aspirated from all wells, replaced with 100 μL MEM + 2% FBS, and cultured for 72 hours further.

For immunoblotting, A549 ACE2 TMPRSS2 cell monolayers were washed 3X with PBS and collected with minimal versene. Ten wells of 96 well plate were pooled per independent experiment for both SARS-CoV-2-infected and uninfected conditions. Pooled cell suspensions were centrifuged (500 × g, 5 mins, room temperature), supernatant was discarded, and cell pellets were lysed for 15 minutes on ice with 100 μL RIPA buffer supplemented with 1 mM PMSF and Halt protease and phosphatase inhibitor cocktail at the manufacturer’s recommended concentration. Cell lysates were clarified (15000 × g, 15 min, room temperature), transferred to a new Eppendorf tube and stored at -80°C. For RNA isolation, monolayers were washed with PBS and 200 μL of TRIzol added to each well of the 96 well plate. Five wells were pooled per independent experiment for both SARS-CoV-2-infected and uninfected conditions into an Eppendorf tube (1 mL TRIzol) followed by the addition of chloroform (200 μL) and centrifugation (12000 × g, 6 min, room temperature). The aqueous layer was transferred to a new Eppendorf with 1 mL ethanol and placed at -80°C for 20 minutes. After centrifugation (12000 × g, 25 mins, 4°C), the supernatant was discarded and RNA pellet was left to air dry for ∼10 minutes and subsequently resuspended in 25 μL RNase free ddH_2_O prior to storage at -80°C.

### Flow Cytometry

HLA-DR and CD86 expression on the surface of moDCs was measured following 72 hrs transduction with NSP5-ZsGreen or empty-ZsGreen pseudotyped lentivirus and subsequent 24 hr stimulation with 100 ng/uL IFN-γ and incubated at 37°C/5% CO_2_. After stimulation 3×10^5^ cells per condition were washed with PBS and blocked for 30 min on ice with FcBlock. Cells were stained on ice for 30 min using eFluor670-FVD and conjugated primary antibodies as indicated in Supplemental Table S2. Cells were fixed with 4% PFA in PBS for 15 min then washed with PBS. Expression levels were measured using a FACSCanto (BD), live moDCs were identified based on FVD-eFluor780 viability dye staining and forward scatter and side scatter profiles. Singlets were gated on the forward area scatter and forward height scatter profiles (**Figure S1A-C**). For cell sorting, singlets were gated on the forward area scatter and forward height scatter profiles, then transduced cells were identified by a positive zsGreen signal and this population sorted into the receiving tube. Flow cytometry data were analyzed using FlowJo (v10.8). All antibodies, dyes, and dilutions used for flow cytometry can be found in Supplemental Table S2.

### Immunoprecipitation and Immunoblotting

Prior to lysis, cells were washed 3× with cold PBS. For NSP5-HDAC2 immunoprecipitations, proteins were reversibly cross-linked using the ReCLIP method [108]. Briefly, cells were incubated for 1 hr at room temperature in PBS + 0.5 mM dithiobis[succinimidyl propionate] and 0.5 mM dithio-bismaleimidoethane). This medium was aspirated, and crosslinking quenched by the addition of 5 mM L-cysteine in 20 mM Tris-Cl, pH 7.4 for 10 min at room temperature. All other immunoprecipitations were performed without cross-linking. Cells were suspended in 300 μL of RIPA lysis buffer, and 50 μL pre-washed anti-DYKDDDDK Tag (L5) beads, rotating for 1 hr. Beads were washed with PBS and immunoprecipitated protein eluted using 0.1 M glycine at pH 2.8, then diluted with 2× Laemmli’s buffer with 5% 2-mercaptoethanol, 1 mM PMSF, and Halt protease and phosphatase inhibitor cocktail at the manufacturer’s recommended concentration. For immunoblotting, cells were lysed with 300 μL RIPA buffer supplemented with 1 mM PMSF and Halt protease and phosphatase inhibitor cocktail at the manufacturer’s recommended concentration. Proteins were loaded on a 4–15% gradient SDS-PAGE gels and transferred onto PVDF membrane. The membrane was blocked for 5 minutes with EveryBlot Blocking Buffer (BioRad) or 5% BSA in TBS-T, incubated overnight at 4°C with the desired primary antibodies (Supplemental Table 2), washed 3 × 5 min with TBS-T, incubated with appropriate IR700 or IR800 secondary antibodies, 1:2,500 dilution, for 1 hr at room temperature in TBS-T. The blots were washed 3 × 15 min washes in TBS-T and visualized with an Odyssey CLx (LI-COR Biosciences, Lincoln, Nebraska). Densitometry was performed in ImageJ/FIJI [109,110].

### Half-Life Determination

The half-life of NSP5 was determined using our established method [111]. Briefly, HeLa cells were split into 6-well plates at a rate of 1.5 × 10^6^ cells/well and transfected with 1 μg/well of wild-type, H41A, C145S, Δ1-192, or Δ199-306 NSP5 using GenJet as per the manufactures instructions. Thirty-six hours later the cells were suspended by trypsinization, counted with a hemocytometer, and 2 × 10^5^ cells/well placed into 6 wells of a 24-well plate. 24 hours later protein synthesis was halted by addition of 50 μg/ml cycloheximide and the cells lysed at 0, 30, 60, 90, 120, and 240 min afterwards. 20 μL of the lysates were immunoblotted with an anti-FLAG antibody as described above, and the quantity of each NSP5 construct determined at each time point using densitometry. Density was then normalized for each construct to the density of that construct at the 0 min timepoint.

### RT-qPCR

Total RNA was isolated from FACS sorted cells transduced with either empty or NSP5-expressing lentiviral vectors using RNeasy Mini Kit as per manufacturer’s instructions. Samples were eluted in 30-50 μL of RNAse-free water. RNA concentration and quality were measured using a NanoDrop 1000 Spectrophotometer. cDNA was obtained from total RNA using the iScript Select cDNA Synthesis Kit according to manufacturer’s instructions using an equal amount of starting RNA and equal mix of the oligo (dT)_20_ primer mixes. RT-qPCR was performed using SsoFast EvaGreen Supermix with an equal amount of starting cDNA. Reactions were run on a QuantStudio 3 Real-Time PCR System for 40 amplification cycles. Relative expression of genes of interest was calculated using the ΔΔCt method, with GAPDH serving as the reference gene.

### Dual-Luciferase Promoter Activity Assay

The MHC II HLA-DRA promoter, the CIITA PI promoter, and the CIITA PIV promoter (**Figure S1D**) were cloned from DNA purified from a human cheek swab using Instagene as per the manufacturer’s instructions using the primers in **Supplemental Table 1** and Phusion DNA polymerase and cloned into pGL4.20 digested with EcoRV using Gibson assembly as per the manufacturer’s instructions. RAW264.7 cells were seeded at 1 × 10^6^ cells/well in a 12-well plate and transfected as indicated with Luc-HLA-DRA, Luc-CIITA pI, Luc-CIITA pIII/IV, Renilla luciferase internal control vector pRL-TK, and NSP5-FLAG or pLVX-IRES-ZsGreen using the Neon electroporation system according to the manufacturer’s protocol. 72 hr post-transfection, cells were lysed with 1x Passive lysis buffer supplemented with EDTA-free HALT protease inhibitor at the manufacturer’s recommended concentration. Dual luciferase assays were performed using a Dual-Luciferase Reporter Assay Kit according to the manufacturer’s instructions, with measurements performed on a Cytation 5 luminescence microplate reader. Firefly luciferase readings were relative to Renilla luciferase readings to account for differences in transfection efficiencies and cell count between samples.

### Immunofluorescence Microscopy

Cells of interest were seeded at a density of 1000 cells/mm^2^ into either 18 mm circular coverslips placed into the wells of a 12-well plate, or into the wells of a custom-printed 15-well imaging chamber [112]. Cells were fixed in 4% paraformaldehyde in PBS for 20 min at room temperature. For plasma membrane staining, cells were stained with 5 μg/mL Alexa Fluor 647-conjugated wheat germ agglutinin for 10 min at 10°C, then fixed in 4% paraformaldehyde in PEM buffer (80 mM PIPES, 1 mM EGTA, 1 mM MgCl_2_) for 10 min at 37°C. If permeabilization was required, fixed cells were treated with permeabilization buffer (PBS + 0.1% triton X-100 + 2.5% BSA); otherwise, cells were blocked with antibody buffer (2.5% BSA in PBS). Anti-FLAG, -MHC II, or -acetyl-lysine were diluted to the concentration indicated in Supplemental Table S1 in antibody buffer and incubated with the cells for 1 hr. Cells were then washed 3 × 15 min with PBS, and then an appropriate secondary antibody added at a 1:1,000 to 1:2,500 dilution in antibody buffer for 1 hr, followed by washing 3 × 15 min with PBS. Samples were either imaged immediately or mounted on a slide using Permafluor before imaging. All incubations and washes were performed at room temperature.

All samples were imaged on a Leica DMI6000B equipped with a Hamamatsu ORCA-Flash4 CMOS camera, fast filter wheels equipped with a Chroma Sedat Quad and custom Fluorescent Resonance Energy Transfer (FRET) filter wheels, operated using Leica LAS-X software. Unless otherwise noted, all cells were imaged using a 100×/1.40 NA objective lens, with Z-stacks acquired with 0.4 μm between slices. Z-stacks were deconvolved in LAS-X using a 10-iteration blinded deconvolution. Images were exported to ImageJ/FIJI for analysis [109,110]. For co-localization studies, the JaCoP plugin was used to calculate the Manders Ratio of NSP5-FLAG and nuclear DAPI or Hoechst staining, or transgene-delineated ER or Golgi markers [113]. To calculate the fraction of NSP5 in the nucleus, a manual region of interest (ROI) was drawn around the nucleus and whole cell and the integrated intensity of each was measured. The fraction of NSP5 in the nucleus was calculated as 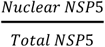. A minimum of 30 transfected cells were quantified per condition for the colocalization and nuclear ratio assays. FRET and FISH-FRET were quantified as described below. To calculate the mean fluorescence intensity of MHC-II in transduced macrophages, the background subtracted channel was thresholded to create a binary mask using the default setting in ImageJ, then a sum slices Z-projection was created. A manual ROI was drawn around the whole cell and the integrated intensity was measured. To determine the surface to cytosol ratio of MHC II in transduced macrophages, the background subtracted channels for wheat germ agglutinin and MHC II were thresholded as described above. Then, the image calculator was used to display all overlapping and non-overlapping pixels, representing MHC II on the cell surface and cytosol, respectively. A summed Z-projection was created, and a manual ROI was drawn around the whole cell to measure the integrated density for both surface- and cytosolic-MHC II. The surface-to-cytosol ratio was calculated as 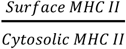 and the fraction of total MHC II on the membrane was calculated as 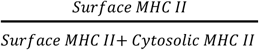.

### Intramolecular FRET

DNA comprised of the human HDAC2 gene with flanking BglII and BamHI restriction sites was synthesized and cloned into pmVenus (L68V)-mTurquoise2 (AddGene #60493) such that a fusion protein of mVenus-HDAC2-mTurqoise2 was produced. To measure NSP5 proteolysis, a codon-optimized DNA sequence encoding the 20 amino acids of the region of the SARS-CoV-2 ORF1ab transcript that codes for the terminal 10 amino acids of NSP4 and initial 10 amino acids of NSP5 (QTSITSAVLQSGFRKMAFPS) was cloned into the BamHI site of the pmVenus (L68V)-mTurquoise2 vector. This site is a known substrate for NSP5 [114]. HeLa cells were then transfected with these constructs with or without NSP5, with mTurquoise2 alone (donor-only sample), with mVenus alone (acceptor-only sample), or with the pmVenus (L68V)-mTurquoise2 vector (positive control). After 24 hours of expression, protein translation was inhibited with 100 μM cycloheximide and the cells incubated for 6 hours to allow NSP5 proteolytic activity to occur in the absence of new protein synthesis. Tiled images of each well were collected, acquiring the donor (mTurqoise2), acceptor (mVenus) and FRET channels at 40× magnification, using the same excitation and camera settings across all samples. FRET efficiency was then calculated using an implementation of the approach of van Rheenen *et al*. [115] using a custom-written script in FIJI. In each repeat, the correction values for donor cross-talk (β, donor-only Ida/Idd), donor cross-excitation (α, acceptor-only Idd/Iaa), acceptor cross-excitation (γ, acceptor-only Ida/Iaa), and FRET cross-talk (δ, acceptor-only Idd/Ida), were calculated using donor-only or acceptor-only images and custom-written scripts in FIJI. FRET efficiency (E_A_) was then calculated in background subtracted images using the formula:

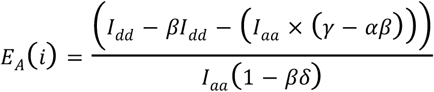

In ImageJ/FIJI the acceptor-only image was then thresholded, and the “Analyze particles” feature used to generate separate ROIs for each cell in each image, and these ROIs were used to quantify the FRET signal of each cell. The maximum theoretical FRET efficiency for the mTurquise2/mVenus FRET pair is 0.3744 [116].

### FISH-FRET

To measure levels of acetyl-lysine at the MHC II and CIITA promoters, human DNA was purified from a cheek swab using Instagene as per the manufacturer’s instructions. 3500 bp amplicons starting before the promoter and ending at the end of the first exon were amplified with Phusion DNA polymerase as per the manufacturer’s instructions using the primers from Supplemental Table S1 (**Figure S1D**). Amplicons were gel purified and cloned into EcoRV-digested pBluescript II using a HiFi Assembly Kit as per the manufacturer’s instructions. The resulting plasmids were labeled with ATTO647N and fragmented using a ATTO647N NT Labeling Kit, producing fragments averaging 200 nucleotides. Primary human macrophages or A549 cells were plated into the 7.5 mm wells of a customized imaging chamber [112], transduced with lentiviral vectors (macrophages) or transfected (A549 cells) with NSP5 or an empty vector, and treated with siRNA as described above. These cells were stained for immune-FISH as per the protocol of Ye *et al*. [117], including wells which were left unstained, or stained only with the donor (acetyl-lysine-Cy3) or acceptor (FISH probes). Briefly, cells were fixed, permeabilized and immunostained for acetyl-lysine as described above. After labeling a secondary fixation was performed for 10 min with 2% PFA. FISH probes were diluted 1:2,500 in hybridization solution (50% formamide, 10% dextran sulfate, 0.3 M NaCl, 30 mM sodium citrate) and denatured at 75°C for 10 min, and then cooled to 37°C. Simultaneously, the cells were incubated at 70°C for 2 min in 70% formamide, 0.3 M NaCl, and 30 mM sodium citrate. The cells were dehydrated by immersing in 75%, 90% and 100% ethanol, 2 min/immersion, then air-dried. The cells were incubated with the denatured FISH probes overnight at 37°C, washed 3 × 5 min with 50% formamide, 0.3 M NaCl, and 30 mM sodium citrate at 42°C, then washed 3 × 5 min with 0.05% Tween 20 in 0.6 M NaCl, and 60 mM sodium citrate. The cells were then washed 3 × 5 min in PBS and immediately imaged.

Tiled images of each well were collected, acquiring the zsGreen, donor, acceptor, and FRET channels at 40× magnification, using the same excitation and camera settings across all samples. FRET efficiency was then calculated as described above. A trained algorithm in Ilastik [118] was used to identify cells based on the acetyl-lysine straining and to classify each cell as zsGreen^+^ (transduced) or zsGreen^-^ (untransduced), collecting a minimum of 500 FISH-labeled loci were analyzed in each experiment. The resulting classifications were exported to FIJI where they were used to assign each FISH probe in the image and the corresponding FRET signal to transduced or untransduced groups. The FRET signal in each sample was then normalized to that observed in the scrambled siRNA-treated, zsGreen^-^ nuclei. The maximum theoretical FRET efficiency of the Cy3/ATTO647N FRET pair is 0.3063 [116].

### NSP5 NLS Analysis

The protein sequence of NSP5 was analyzed for the presence of monotonic and bipartite nuclear localization signals using the default settings on four different prediction algorithms: cNLS Mapper (https://nls-mapper.iab.keio.ac.jp/cgi-bin/NLS_Mapper_form.cgi), 4 state HMM on NLStradamus (http://www.moseslab.csb.utoronto.ca/NLStradamus/), seqNLS (http://mleg.cse.sc.edu/seqNLS/), and in InterProScan (https://www.ebi.ac.uk/interpro/search/sequence/).

### Phylogenetic Analysis

Using the protein sequence of human HDAC2 and the NCBI BLASTp tool (https://blast.ncbi.nlm.nih.gov), the protein sequences of HDAC2 from a range of species representing the major vertebrate clades were identified. The same approach, using the protein sequence of NSP5 from SARS-CoV-2 was used to identify NSP5 protein sequences across the four coronavirus genera. These sequences were imported into MEGA XI, and a MUSCLE alignment of the protein sequences was generated [119]. Pairwise distances were then calculated using a Poisson model assuming uniform rates across sites, and maximum likelihood trees were generated using a 500-iteration bootstrapping approach. Per-residue conservation was quantified using the Shannon Entropy calculator on the Protein Residue Conservation Prediction server (https://compbio.cs.princeton.edu/conservation/) [120].

### Statistical Analysis

Using GraphPad Prism, a Shapiro-Wilk test was used to determine whether data was parametrically or non-parametrically distributed, and data was then analyzed using an appropriate 2-tailed statistical test, as indicated in the figure legends. Parametric data is presented as mean ± SEM, while non-parametric data is presented as box-and-whisker or violin plots with median and quartiles.

## Preprint Notice

This publication is a scientific preprint and has not undergone peer review.

## Data Availability

All data except for raw (unprocessed and unanalyzed) microscopy images are contained within the manuscript. These images are available from the corresponding author upon request, Bryan Heit.

## Supplemental Figures

This article contains Supplemental Figures.

## Acknowledgments

We would like to thank Kristin Chadwick from the London Regional Flow Cytometry Facility for her assistance in the design and optimization of the flow cytometry assay.

## Funding and Additional Information

This work was funded by a Canadian Institutes of Health Research (CIHR) Project Grant (PJT-162203) to BH. AL, BHD, and ENB were funded by Canada Graduate Scholarships – Master’s (CGSM) from the CIHR. AA was funded by AA funded by an Ontario Graduate Scholarship and RGE Murray Scholarship. The funding agencies had no role in study design, data collection and analysis, decision to publish, or preparation of the manuscript.

## Author Contributions

Nima Taefehshokr: Conceptualization, Methodology, Formal analysis, Investigation, Writing – Original Draft, Writing – Review and Editing. Alex Lac: Conceptualization, Methodology, Formal analysis, Investigation, Writing – Original Draft, Writing – Review and Editing. Angela M Vrieze: Conceptualization, Formal analysis, Validation. Brandon H Dickson: Investigation, Formal analysis. Catherine Jung: Investigation, Formal analysis. Amena Aktar: Investigation, Writing – Review and Editing. Eoin N Blyth: Methodology and Validation. Corby Fink: Investigation, Formal analysis. Jimmy D Dikeakos: Conceptualization, Supervision, Funding acquisition. Gregory A Dekaban: Conceptualization, Supervision, Funding acquisition. Bryan Heit: Conceptualization, Methodology, Formal analysis, Investigation, Writing – Original Draft, Writing – Review and Editing, Supervision, Funding acquisition, Project administration.

