## Supplemental Tables 1 & 2, Supplemental Figures S1 - S5 for "SARS-COV-2 NSP5 Antagonizes MHC II Expresion by Subverting Histone Deacetylase 2"

### Supplemental Materials

**Table S1: PCR Primers Used in This Study**

| Product | Forward Primer | Reverse Primer |
|---|---|---|
| MHC II FISH Probe | ACGGT ATCGA TAAGC TTGAT TTTAT TCTGA TAGGG ATCTA TTCCA | CCGGG CTGCA GGAAT TCGAT CTCAG CACCT ACCTT TGATA |
| CIITA pI FISH Probe | ACGGT ATCGA TAAGC GGCGT GAACC CAGGA GGC | CCGGG CTGCA GGAAT TCGAT ACCTT GGGGC TCTGA CAGGT A |
| CIITA pIII/IV FISH Probe | ACGGT ATCGA TAAGC TAAAA AGGCC GGGAA AGCAT CTTAA TTTAG CGTG | CCGGG CTGCA GGAAT TCCTG TGGAG CAACC AAGCA CCTAC T |
| MHC II luciferase reporter | AGCTC GCTAG CCTCG AGGAT TCCGT GATTG ACTAA CAGTC | CGCCG AGGCC AGATC TTGAT GAATA AAAGA AAAGA GAATG TGGG |
| CIITA luciferase reporter | AGCTC GCTAG CCTCG AGGAT GATAT TGGCA GCTGG CACCA | CGCCG AGGCC AGATC TTGAT CAGCT CAGAA GCACA CAGCC |
| RFX5 RT-qPCR | TCCTT CAGTT CCATC GTTGA G | TTCAG CTGTC CTCTT GACAC C |
| CIITA RT-qPCR | CTGAA GGATG TGGAA GACCT GGGAA AG | ACCCT CGTCC CCGAT CTTGT TCTCA CTC |
| MHC II RT-qPCR | CGAGT TCTAT CTGAA TCCTG | GTTCT GCTGC ATTGC TTTTG C |
| GAPDH RT-qPCR | TCAAG GCTGA GAACG GGAAG | CGCCC CACTT GATTT TGGAG |
| NSP5 H41A | P-CTGTG ATCTG CACCT CTGAA GACAT GC | P-CTCTT GGACA GTAAA CTACG TCATC AAGCC A |
| NSP5 C145S | P-CTGGT AGTGT TGGTT TTAAC ATAGA TTATG ACTGT GT | P-ATGAA CCATT AAGGA ATGAA CCCTT AATAG TGAAA TTG |
| NSP5$^{\Delta 1\text{-}192}$ | P-GCAGC TGGTA CGGAC ACAAC TATTA C | P-CATGA ATTCA CCGGA AATAG ATCCT CTAGT AGAG |
| NSP5 $^{\Delta 199\text{-}306}$ | P-GCATC ACCGG TAGAC TACAA GGACC | P-TGTGT CCGTA CCAGC TGCTT G |

*Note: "P-" indicates that the 5' end of the primer is phosphorylated.*

**Table S2: Antibodies and Other Staines Used in This Study.**

| Target | Clone | Species/Isotype | Company | Concentration* |
|---|---|---|---|---|
| PE-CD86 | BU63 | MouseIgG | BioLegend | FC: 0.5 µg/mL |
| APC-HLA-DR | L243 | Mouse IgG | BioLegend | FC: 0.25 µg/mL |
| FVD-eFluor780 | -- | -- | ThermoFisher | FC: 1 µL/mL |
| Human TruStain FcX | -- | -- | BioLegend | FC: 50 µL/mL |
| FLAG | L5 | Rat IgG2 | Sigma-Aldrich | IB: 0.5 µg/mL<br>IP: 1.0 µg/mL |
| V5 Tag | polyclonal | Rabbit | Sigma-Aldrich | IB: 0.5 µg/mL<br>IP: 1.0 µg/mL |
| HDAC2 | HDAC2-62 | Mouse IgG2b | Sigma-Aldrich | IF: 0.5 µg/mL<br>IB: 0.5 µg/mL<br>IP: 1.0 µg/mL |
| MHC II, pan-human | RBM1-2967-P1 | Rabbit IgG | ThermoFisher | FC: 0.5 µg/mL |
| GAPDH | D16H11 | Rabbit IgG | Cell signaling | IB: 1 µg/mL |
| α-tubulin | 236-10501 | Mouse IgG1 | ThermoFisher | IB: |
| Acetyl-Lysine | clone 1C6 | Mouse IgG | Abcam | IF: 1 ng/mL |
| DAPI | -- | -- | ThermoFisher | IF: 0.5 µg/mL |
| Hoechst | -- | -- | ThermoFisher | IF: 1 µg/mL |
| Wheat Germ Agglutinin-Alexa Fluor 647 | -- | -- | ThermoFisher | IF: 5 µg/mL |

** Concentrations used for flow cytometry (FC), immunofluorescence (IF), immunoblotting (IB), or immunoprecipitation (IP).*

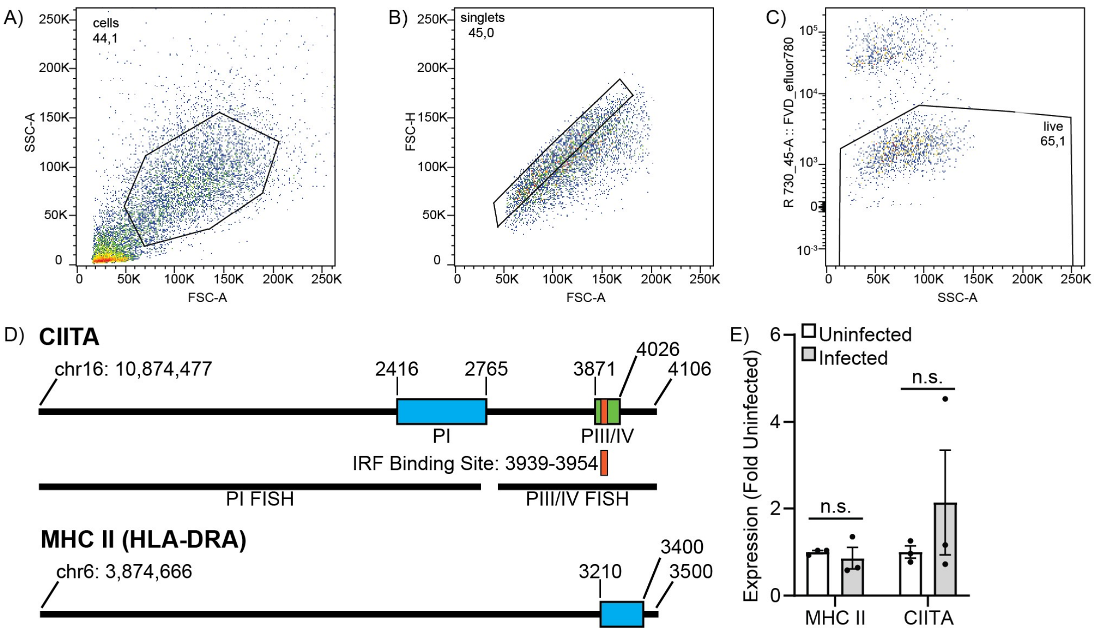


**Supplemental Figure S1. Flow Gating and Cloning Strategy. A-C)** Human moDCs were transduced with lentiviral vectors bearing a zsGreen selectable marker and identified by excluding cell debris using the FSC-A/SSC-A scattergrams (A), selecting selected via FSC-A/FSC-H (B), and live moDCs identified by gating on SSC-A and the cell viability dye eFluor670-FVD (C). Up to 10,000 cells were recorded per condition in each experiment. **D)** Regions cloned for the FISH-FRET and dual-luciferase vectors. For CIITA, two FISH promoters were cloned covering the PI promoter and its 5' region (PI FISH), as well as the region between the end of the PI promoter and 80 bp downstream of the PIII/PIV promoters (PIII/IV FISH). Dual-luciferase vectors containing the PI (blue), PIII/IV (green), and IRF-binding site (orange) deleted PIII/IV promoters were generated. For MHC II, a FISH probe consisting of a 3500 bp region covering from 3200 bp 5' to the HLA-DRA promoter to 100 bp after the promoter was cloned, as was a dual-luciferase vector containing the promoter itself (blue). **E)** RT-PCR quantification of MHC II and CIITA expression in A549 cells that were uninfected or infected with the USA-WA1/2020 strain of SARS-CoV-2. Data is expressed as ΔΔCt, normalized to uninfected. Data quantifies (E) or is representative of (A-C) 3 independent experiments. n.s. = $p > 0.05$, Mann-Whitney U Test.

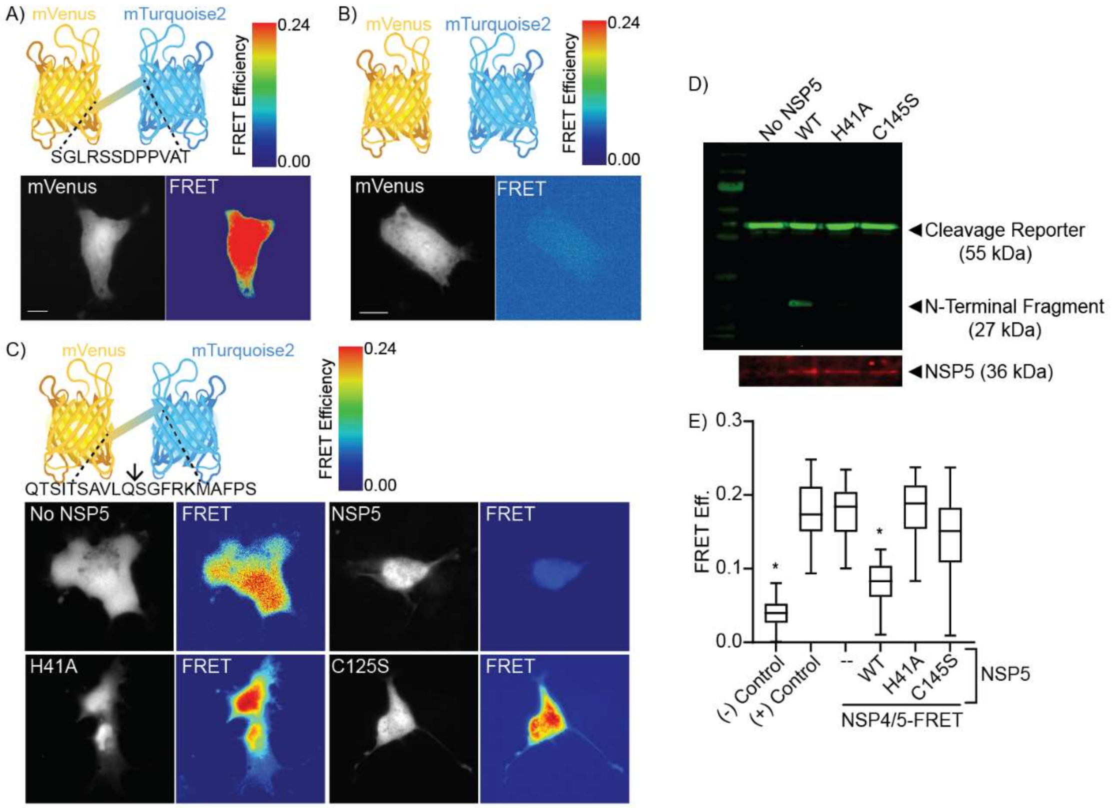


**Supplemental Figure S2. Quantification of NSP5 Proteolytic Activity. A)** Structure (top) and FRET signal (bottom) of the positive FRET control, comprised of mVenus and mTurquoise2 separated by a 12-amino acid linker. **B)** Structure (top) and FRET signal (bottom) of the negative FRET control, comprised of mVenus and mTurquoise2 expressed as separate proteins. **C)** *top:* structure of the NSP5 proteolysis intramolecular FRET probe in which mVenus and mTurquoise2 are linked by the final 10 amino acids of SARS-CoV-2 NSP4 and the first 10 amino acids of SARS-CoV-2 NSP5. The known NSP5 cleavage site is indicated by the arrow. *Bottom:* localization and FRET efficiency of the NSP5 proteolysis probe when expressed in the absence of NSP5 (No NSP5), in the presence of wild-type NSP5 (NSP5), or when expressed with the $NSP5^{H41A}$, or $NSP5^{C145S}$ point mutants. **D)** Quantification of NSP5 proteolysis reporter cleavage by immunoblotting for mVenus showing the expected mass of the uncleaved reporter (55 kDa) and the N-terminal fragment produced by cleavage (27 kDa). **E)** Measurement of NSP5 proteolysis reporter cleavage via quantification of whole-cell FRET. Cells are transfected with the positive or negative control vectors, or with the NSP5 cleavage reporter plus either empty vector (--), wild-type NSP5 (WT), or the $NSP5^{H41A}$ or $NSP5^{C145S}$ point mutants. n = 5, * = $p < 0.05$ compared to the positive control (+) Control vector, Kruskal-Wallis test with Dunn correction.

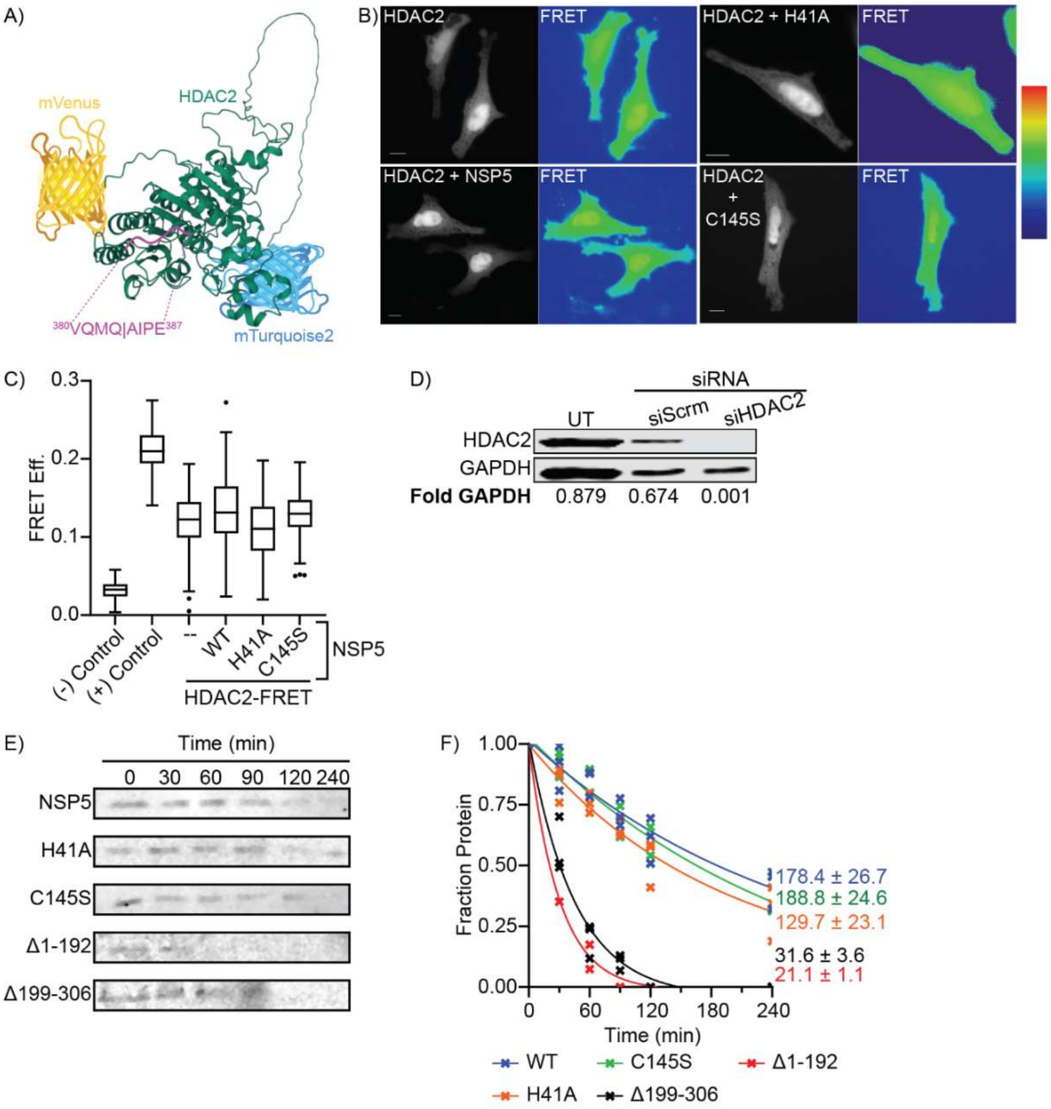


**Supplemental Figure S3: HDAC2 Cleavage and NSP5 Half-Life. A-B)** Structure (A) and FRET signal (B) of a HDAC2 intramolecular cleavage FRET probe in which mVenus is fused to the N-terminus, and mTurquoise2 to the C-terminus, of human HDAC2. The putative NSP5 proteolytic site is indicated in purple. **C)** Quantification of HDAC2 cleavage by NSP5 as quantified by whole-cell FRET. Cells are transfected with either the FRET negative control [(-) Control], FRET positive control [(+) Control], or with the HDAC2-FRET reporter plus one of empty NSP5 vector (-), wild-type NSP5 (WT) or with the H41A or C145S NSP5 mutants. **D)** Confirmation of HDAC2 knockdown in cells that are untreated (UT), transfected with a cell-permeant non-targeting siRNA (siScrm), or transfected with a cell-permeant HDAC2-targeting siRNA (siHDAC2). **E)** NSP5 lifespan immunoblots from cells transfected with wild-type NSP5 (NSP5), the $NSP5^{H41A}$ or $NSP5^{C145S}$ point mutants, or the $NSP5^{\Delta 1\text{-}192}$ and $NSP5^{\Delta 199\text{-}306}$ deletion mutants. Protein synthesis was inhibited at t= 0 min with cycloheximide and equal volumes of cell lysate loaded from each indicated timepoint. **F)** Determination of NSP5 half-life by densitometry, with non-linear regression used to calculate the half-life. * = $p < 0.05$ compared to the positive control (+) Control vector, Kruskal-Wallis test with Dunn correction.

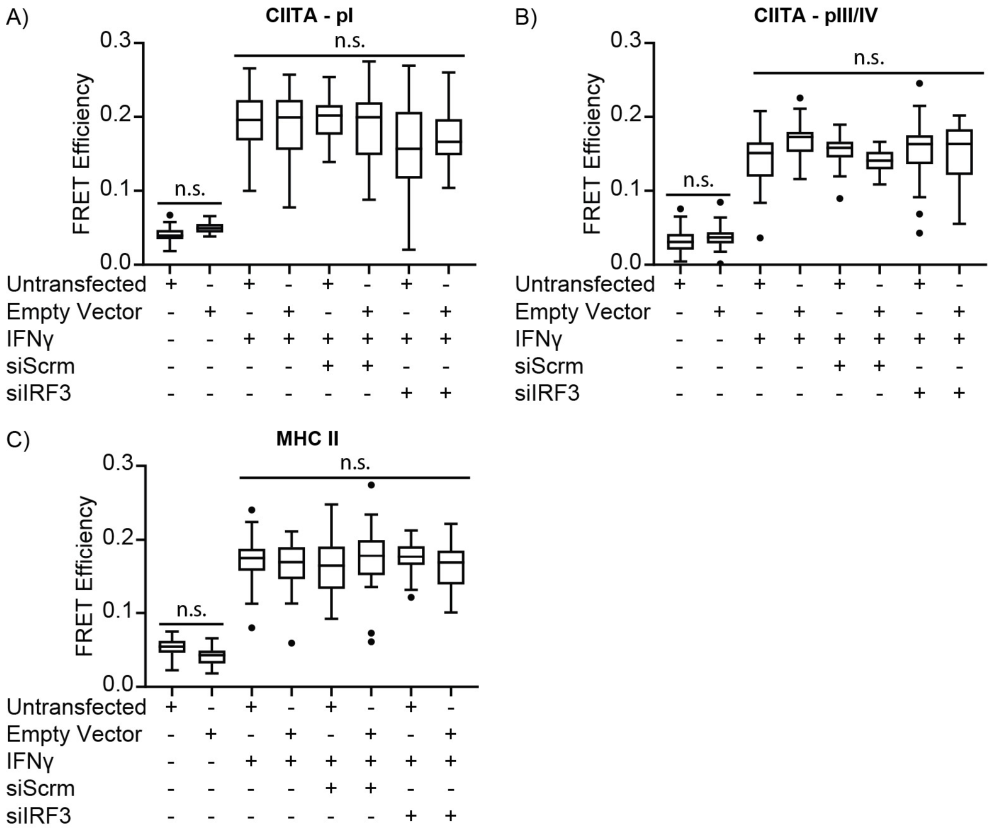


**Supplemental Figure S4: Control FISH-FRET Data for IRF3 siRNA Knockdown.** Quantification of FISH-FRET at the CIITA pI (A), CIITA pII/IV (B) and MHC II (C) promoters in A549 cells co-transfected with empty vector in lieu of NSP5-expression vectors, and then treated with IFN-γ and a non-targeting (Scrm) or IRF3-depleating siRNA. Data is presented as quartiles, n = 3, * = $p < 0.05$; n.s. = $p > 0.05$ compared to Empty (B), WT (D), or the indicated groups (E-G), Kruskal-Wallis test with Dunn correction.

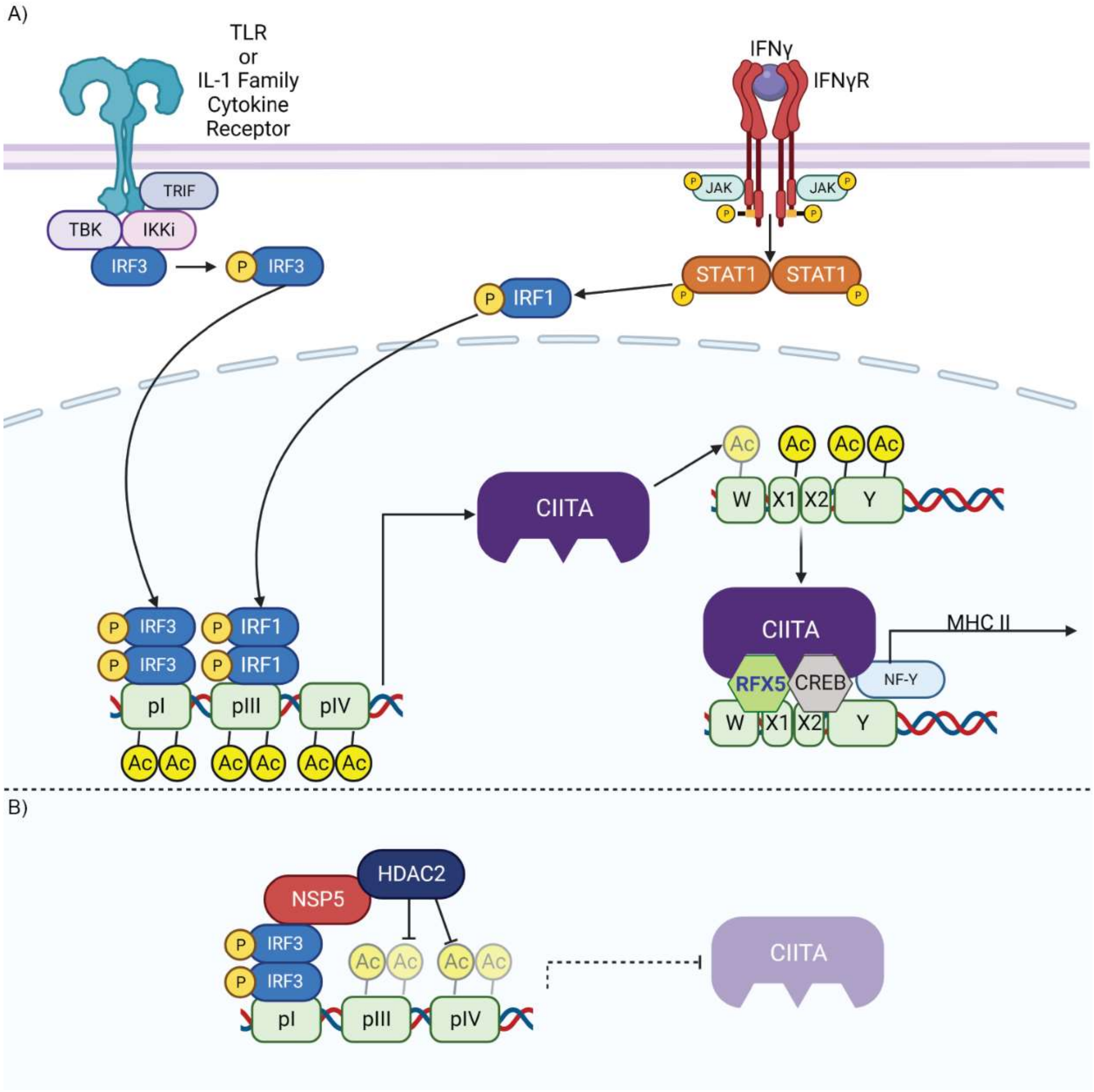


**Supplemental Figure S5: Model of MHC II Transcriptional Control and NSP5 Activity at the MHC II Promoter. A)** MHC II expression in myeloid cells is induced by a combination of IFN-γ/STAT1 and TLR or IL-1 family cytokine signaling, which respectively activate IRF1 and IRF3. IRF1 and IRF3 then bind to the CIITA promoter and induce CIITA expression (*left*). Once synthesized, CIITA directly acetylates histones in the MHC II promoter via its intrinsic acetyltransferase activity. Once the MHC II promoter is acetylated, CIITA, RFX5, and NF-Y form an activating complex on the W/X1/X2/Y motifs found in the core of the MHC II promoter, inducing expression of MHC II (*right*). **B)** During SARS-CoV-2 infection, NSP5 binds to HDAC2, and via interactions with IRF3, delivers HDAC2 to the CIITA promoter. Here, HDAC2 deacetylates and inactivates the CIITA promoter, thereby suppressing expression of CIITA, with the loss of CIITA expression then leading to the cessation of MHC II expression. Figure produced in BioRender.
